# Histologically resolved spatial multi-omics of human oral squamous cell carcinoma

**DOI:** 10.1101/121616

**Authors:** Tao Chen, Chen Cao, Jianyun Zhang, Aaron Streets, Yanyi Huang, Tiejun Li

## Abstract

Both the composition of cell types and their spatial distribution in a tissue play a critical role in cellular function, organ development, and disease progression. For example, intratumor heterogeneity and the distribution of transcriptional and genetic events in single cells drive the genesis and development of cancer. However, it can be challenging to fully characterize the molecular profile of cells in a tissue with high spatial resolution because microscopy has limited ability to extract comprehensive genomic information, and the spatial resolution of genomic techniques tends to be limited by dissection. There is a growing need for tools that can be used to explore the relationship between histological features, gene expression patterns, and spatially correlated genomic alterations in healthy and diseased tissue samples. Here, we present a technique that combines label-free histology with spatially resolved multi-omics in un-fixed and unstained tissue sections. This approach leverages stimulated Raman scattering microscopy to provide chemical contrast that reveals histological tissue architecture, allowing for high-resolution *in situ* laser micro-dissection of regions of interests. These micro-tissue samples are then processed for DNA and RNA sequencing to identify unique genetic profiles that correspond to distinct anatomical regions. We demonstrate the capabilities of this technique by mapping gene expression and copy number alterations to histologically defined regions in human squamous cell carcinoma (OSCC). Our approach provides complementary insights in tumorigenesis and offers an integrative tool for macroscale cancer tissues with spatial multi-omics assessments.

## Introduction

Recent technological developments for single-cell genomic analysis have led to high-resolution characterization of cellular heterogeneity in tissues and organs [1-2]. Most of these techniques require the dissociation of millimeter-scale biological samples, or larger, before the extraction of genomic material, limiting the spatial resolution of these techniques to the size of the dissected tissue sections [3-4]. While high-throughput single-cell RNA sequencing has been a powerful tool for classifying cell types and revealing cell states, cellular state and function are strongly influenced by the cell’s microenvironment, including chemical and physical context as well as the composition of neighboring cells [5-6]. Therefore, comprehensive cellular characterization of biological systems would ideally leverage the combination of high-throughput molecular analysis and high spatial resolution. Laser capture microdissection (LCM) [7-9] enables the recovery of sub-millimeter regions of interest from tissue, allowing for microscopy to guide the selection of cells based on the morphology of the microenvironment. In this way, LCM links histology with molecular measurements, including genome and transcriptome-wide profiling with high-throughput sequencing. However, LCM typically requires fixation and staining to generate histological contrast, and these steps can perturb or even degrade the nucleic acid composition of cells [10-11]. Furthermore, the contrast used to guide dissection in these applications is limited by the chosen staining method [12].

Here we present an integrative approach for mapping molecular and cellular heterogeneity in tissue samples that leverages stimulated Raman scattering (SRS) microscopy [13] to generate histological images of fresh or frozen tissue slices followed by *in situ* micro-dissection and multi-omic sequencing analyses. This technique, called SRS micro-dissection and sequencing (SMD-seq), images tissue sections without staining, and instead provides contrast based on the intrinsic chemical composition of the biological specimen. SRS generates signal from molecular vibrations that are specific to chemical bond composition. Imaging a biological sample at multiple Raman frequencies can therefore provide composite images with morphological contrast that corresponds to chemical variation [14-17]. After label-free imaging of fresh or frozen tissue slices, regions of interest are dissected directly with the SRS excitation laser, and immediately processed enabling efficient recovery of RNA and DNA from small numbers of cells within a region of interest (Fig 1a, materials and methods). The combination of quantitative chemical imaging and high-resolution genomic analysis facilitates multimodal characterization of the cellular heterogeneity in complex tissues.

**Fig-1.**
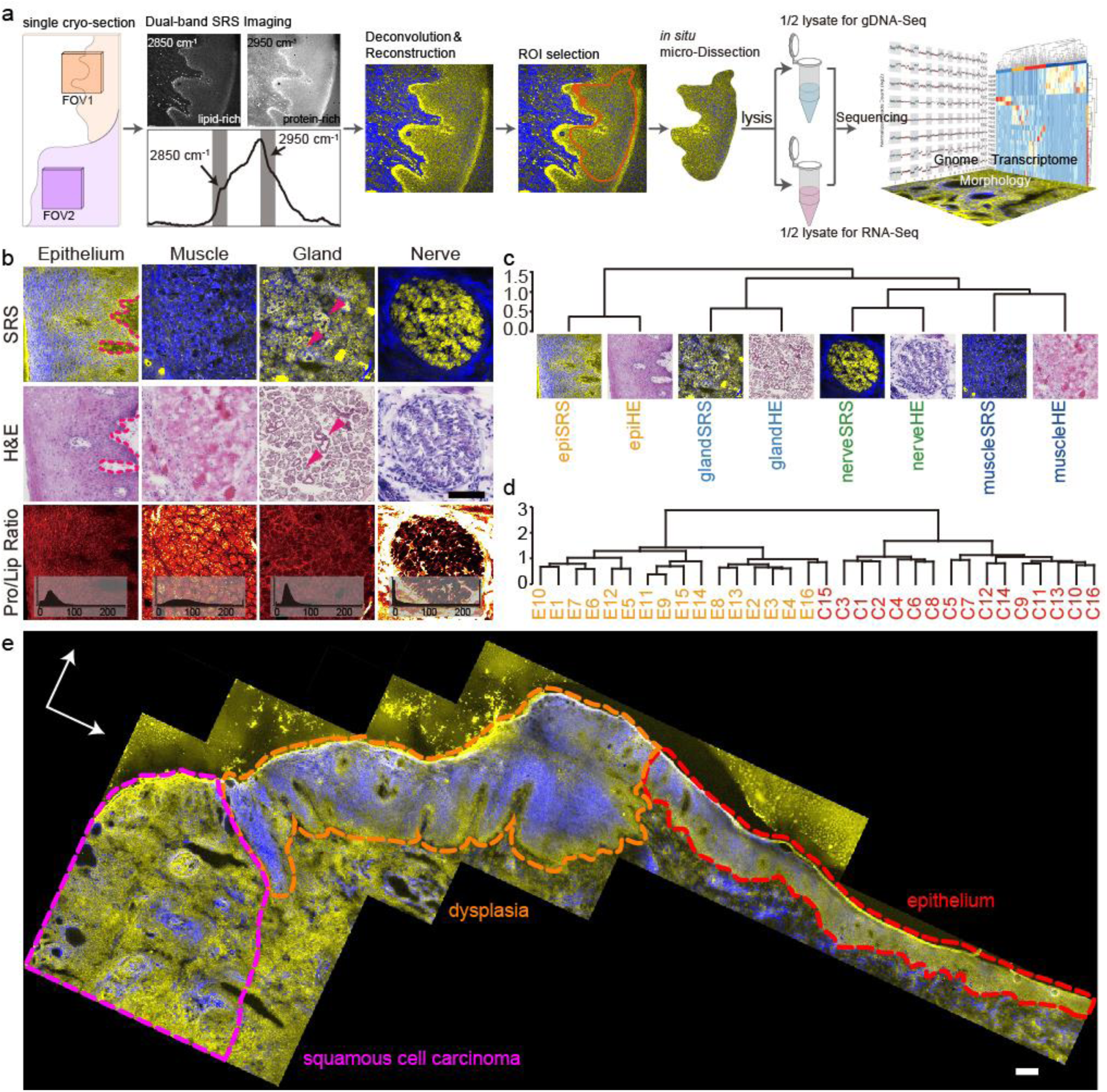
Principle of SMD-seq and evaluation of SRS imaging. a) the pipeline of SMD-seq, inset of FOV2 represents the complexity around single cancer nest. b) the comparison between SRS and H&E images of different tissues, with the PLR images of SRS. c) unsupervised clustering of HOG features of SRS and H&E images from different tissue types. d) unsupervised clustering of SRS images of epithelium and cancer. e) stiched SRS image of a whole sample, showing revealed morphologies including cancer, dysplasia, and epithelium. The magenta stars indicate the keratin pearls. Scale bars are 200 μm.

The ability to map molecular and cellular heterogeneity is particularly important in cancer. Heterogeneity in cancer is expressed in many facets, including the evolving genetic changes in distinct subpopulations [18-19], and the progressive abnormalities in morphology of the cells, typically identified by histopathology [20]. Molecular and phenotypic aberrant variations are not only common between tumors of different tissue and cell types, but also within a tumor derived from the same patient or even from the same tissue [21]. The clonal diversity during tumorigenesis, in the context of histology and genetic divergence, influences each individual tumor’s diagnosis and response to treatment [22]. It is therefore important to characterize the tumor heterogeneity in a comprehensive and precise way.

In this study, we demonstrate the utility of SMD-seq by spatially mapping genomic and transcriptomic profiles in human oral squamous cell carcinoma (OSCC). OSCC is a major subtype of head and neck squamous cell carcinoma (HNSCC). Its aetiological and biological heterogeneity are influenced by distinct risk factors [23-24], which further lead to high morbidity and relatively low overall 5-year survival rate. OSCC is highly invasive, leading to large variation in cellular phenotypes over small length scales [25-26]. OSCC cancer nests, for example, measuring as small as 100 μm in diameter, are often interlaced with normal tissues composed of non-cancerous cells, and it is challenging to dissect this heterogeneity with existing methods [25]. Additionally, mutation patterns in OSCC have been shown to differ from other subsites of HNSCC due to its specific anatomical location and tissue characteristics [27]. We used SRS histology to identify progressive abnormalities in the morphology of OSCC cancer cells, and in situ laser micro-dissection followed by DNA and RNA sequencing identified unique molecular profiles that correspond to distinct anatomical regions of tissue samples from multiple individuals. Transcriptomic analysis was used to identify gene expression profiles that corresponded to specific tissue types and cancer cells, which corroborated histological analysis. Furthermore, transcriptomic and genomic analysis revealed gene fusion events and copy number variations that delineated cancer progression within tissue sections. This combination of imaging and multi-omic analysis demonstrates how SMD-seq can provide a powerful tool to accurately investigate complex inter- and intra-tumor heterogeneity and has the potential to be scalable to any architecturally complex tissue.

## Results

### Label-free SRS microscopy accurately revealed histological features of OSCC

The recent and rapid development of SRS-based histology [14-17] has proven this technique to be a powerful supplement to traditional histological staining for many tissues. We first examined the ability of SRS imaging to reconstruct histological features in unstained cryo-sections of OSCC that corroborated information obtained with H&E staining of OSCC sections. We generated two-color SRS composite images of 30 μm cryo-sections biopsied from various tissues that provide contrast between lipid-rich and protein-rich structures by exciting CH2 symmetric vibration and CH3 vibrations respectively (methods, Fig. S1). We then compared these reconstructed images to adjacent 5 μm cryo-sections that were sampled from the same biopsies and conventionally stained by H&E. The lipid-protein contrast in the SRS images recovered the characteristic morphological features of each tissue, including the variation of cell shape and base membrane in the epithelium, the epithelial cells forming the duct wall in the gland tissue [28], and the nerve bundles (Fig 1b). In addition to recapitulating morphological information, SRS images provide information about the chemical composition of the tissue samples. For example, both the muscles and the perineurium, a collagen-rich structure [29-30], around nerve revealed high protein content in SRS images (Fig 1b). Variability in the chemical composition of ROIs can be further evaluated by the histogram of the protein-to-lipid ratio (PLR) from different tissue types, reflecting a distinct molecular signature of the cells within the tissue. In order to confirm that the histopathological utility of SRS was not limited to OSCC, we also identified characteristic features of two other oral cavity diseases, the Warthin’s tumor and the Mucoepidermoid Carcinoma (Fig. S2). These results demonstrate the generalizability of SRS imaging for oral cavity histology.

We then quantified the similarity between morphological information retained in SRS and H&E images using the histogram of orientation gradient (HOG) [31]. The HOG describes the texture of an image based on intensity variation, which primarily reflects cellular packing in our images. Unsupervised clustering of the HOG from all eight images presented in figure 1b, successfully clustered images by tissue type, regardless of imaging modality (Fig-1c). This analysis indicates that SRS images recovered similar cellular organization patterns as H&E images. In addition to distinguishing different types of tissue, the ability to distinguish cancerous epithelium from normal epithelium has potential clinical implications. The cellular organization patterns between these tissue states are distinct and can be decoded by HOG features. Unsupervised clustering of the HOG features of 32 SRS images, including 16 cancerous and 16 normal epithelia (Fig. S3), stratified tissue images into two groups, accurately reflecting disease state with only one misidentification. (Fig-1d). Furthermore, the correlation matrix of all 32 samples’ HOG features revealed a weaker correlation among cancers than epithelia (Fig. S4), implying greater variation in cell-packing patterns between individual cancer nest. This heterogeneity could be related to the distinct micro-environment of the cancer nests. For example, the cancer nests in P4 invaded heavily into muscle tissue, while in other patient samples the cancer nests were typically located in connective tissue (Fig. S5). We examined the morphology restoration at different scales on stitched image of a whole cryo-section (9×4 mm ^2^). Comparing with corresponding H&E images, SRS revealed identical features at all scales. Selected regions of interest from SRS images can be localized in H&E images through correlation of their respective HOG (Fig-1e, Fig. S6). The key morphological features were identified in the large-scale SRS image depicted in figure 1e, including normal epithelium, dysplasia, cancer, and those keratin pearls correlated with OSCC differentiation.

### SRS guided micro-dissection enabled high-quality mRNA and DNA sequencing of cancer nests

In the SMD-seq workflow, after an ROI is identified with SRS histology, that region is immediately excised and recovered from the bulk tissue section for downstream genomic analysis. This is achieved by increasing the power of the pump line of the SRS excitation source and scanning the focal spot around a user-defined ROI boundary (Methods). To validate the accuracy of ROI dissection, we performed H&E staining on dissected tissue, as well as adjacent tissue sections (Fig-2a). Because microdissection is performed with the scanning laser, the ROI is defined with high resolution (Fig. S7a). and can be recovered immediately after imaging. Depending on the ROI size and tissue type, the dissected sample typically contained about 230 cells on average (Fig. S7b). SRS imaging and dissection can be performed in rapid succession, allowing the dissection of multiple ROIs from a single tissue section (Fig-2b), therefore all the cancer microtissues can be compared to a corresponding normal microtissue from the same sample as a control.

**Fig-2.**
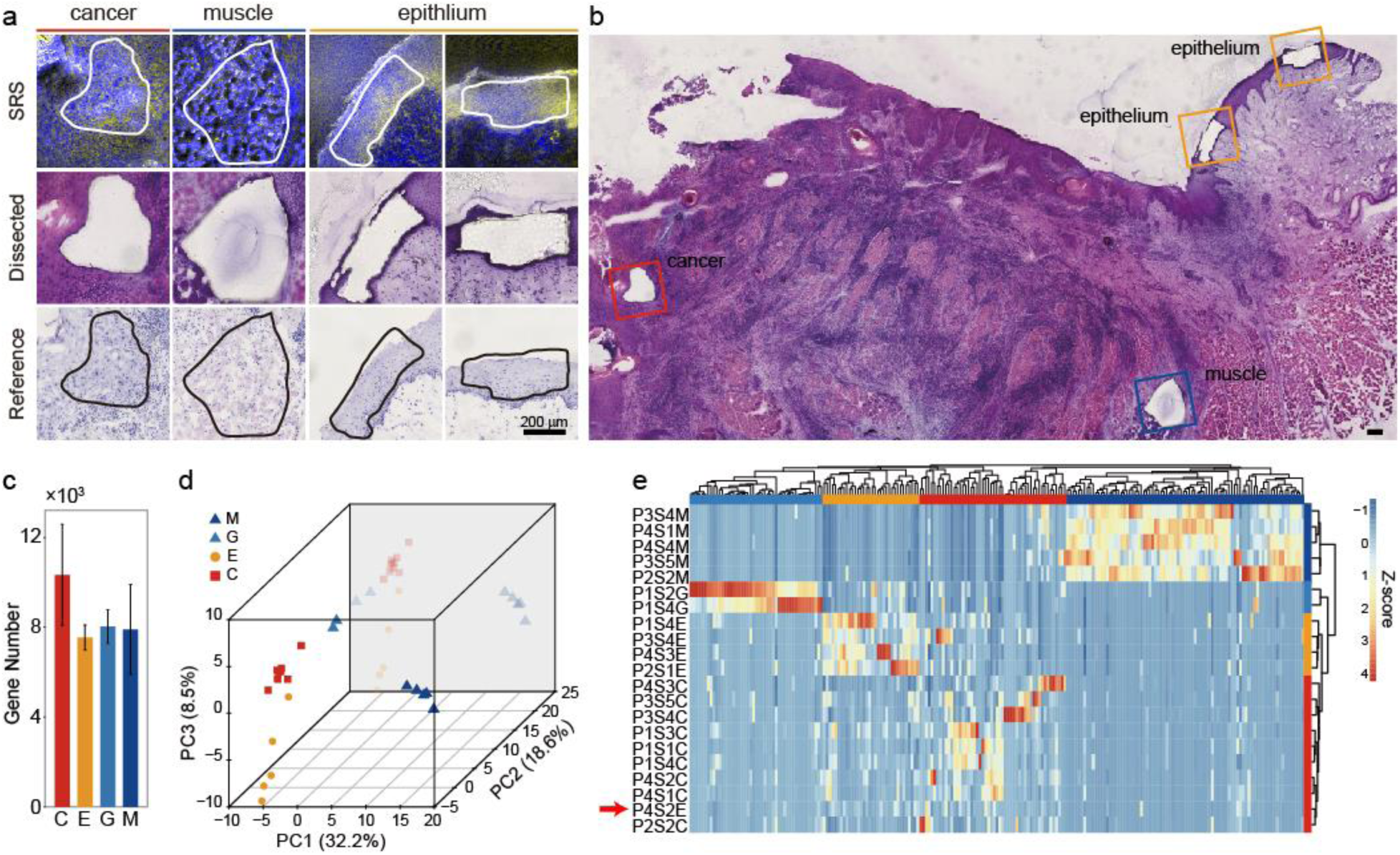
SRS image guided micro-dissection and evaluation of mRNA recovery. a) comparison between SRS images, H&E stained and unstained micro-tissue section. b) epithelium, muscle, and cancer from the same section c) statistics of gene number per sample d) PCA plot of all expressed genes. e) unsupervised clustering of 217 differently expressed genes from micro-dissected samples. Scale bars are 200μm.

After recovery, each micro-sample was lysed and divided into two equal aliquots for parallel DNA and mRNA extraction. One of the primary advantages of SMD-seq, is the efficient recovery of high-quality mRNA, which is made possible because SRS histology avoids fixation, staining or any other chemical perturbations which may degrade RNA or inhibit recovery. Quantification of house-keeping genes showed that unstained cryo-sections can preserve over 20-fold more mRNA than H&E stained sections (Fig. S8). Another advantage of SMD-seq is that high-resolution identification of ROIs allows for more precise selection of tissue and cell types of interest. Comparing with head and neck squamous cell carcinoma samples collected for the Cancer Genome Atlas, (TCGA), the SMD-samples presented significantly less cell type contamination (P value = 0.0098, Fig. S7c-e, Methods).

We applied SMD-Seq to 13 cryo-sections from four patients (3 males, 1 female) who suffered various stages of OSCC at different ages (Methods). We collected 28 in situ micro-dissection samples for sequencing, and 27 samples passed the quality control based on mRNA recovery, for further analysis (Supplementary Table 1-3, Methods). These micro-samples included 12 normal (5 muscle; 2 gland; 5 epithelium) and 9 cancer samples for RNA-Seq, and 13 normal (muscle, n=5; gland, n=2; epithelium, n=6) and 8 cancer samples for DNA-Seq. RNA-seq recovered around 9,000 genes per micro-sample on average (FPKM>0.1, Fig-2c) and highly correlated expression among samples of the same tissue, indicating technical reproducibility (Fig. S9).

Principal component analysis separated micro-samples into four groups of similar gene expression profiles which recapitulated the known tissue types (Fig-2d). Unsupervised hierarchical clustering of micro-samples using the top 217 differently expressed genes (Methods) grouped these clusters as distinct tissue types, three of which were then annotated using a panel of known marker genes for epithelium, gland and muscle in agreement with the corresponding tissue types identified by SRS histology (Fig-2d, Fig. S10). The fourth group, which was consistently identified as cancer by SRS images and, contained highly expressed OSCC marker genes including GSTP1, AKR1B10, FTH1, and FTL (Fig. S11). GSTP1 expression is known to be associated with high malignancy and poor survival rate [32-33]. Expression levels of GSTP1 were further validated with immunofluorescence staining, confirming the confinement of GSTP1 expression to cancer nests (Fig. S12). While the epithelial to mesenchymal transition (EMTs) related gene, such as KRT13 (Fig-3a) [34-35], was expressed at lower levels, indicating an enhanced migratory capacity and invasiveness [36]. These OSCC samples originated from the epithelium, as determined by imaging, but revealed a distinct expression profile compared to the healthy epithelium samples Unexpectedly, P4S2E, which was identified as epithelium by SRS imaging, clustered with the cancer samples based on gene expression profiles (Fig-2e, red arrow). Principal component analysis also indicated that P4S2E, was more similar to the cancer samples than the epithelium samples (Fig-2d). This observation indicates that micro-sample P4S2E may have come from a region of the biopsy in which these epithelial cells were developing into carcinoma. This disagreement between histology and gene expression could only be captured by SMD-seq, where independent morphology and gene expression data are derived from the same sample.

**Fig-3.**
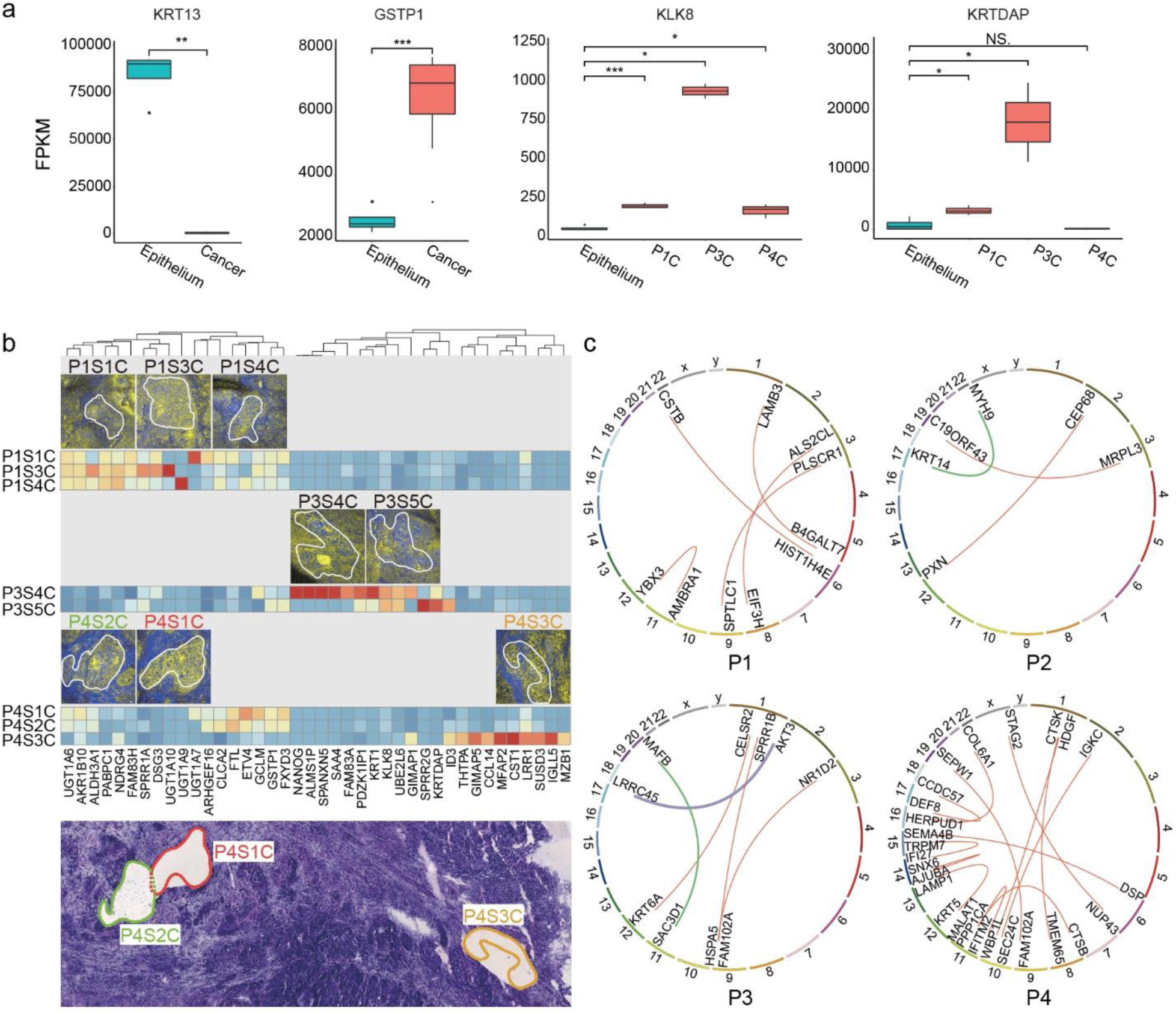
Heterogeneity of gene expression. a) Expression levels of genes in cancer samples among different patients, in comparison with epithelium. b) the inter-patient and intra-patient cancer heterogeneity. The bottom figure is a projection of three slides, showing the relative positions of different cancer nests. c) inter-patient heterogeneity of gene fusion events.

### Molecular Heterogeneity in OSCC

SMD-seq provides the ability to compare the genome and transcriptome of specific tissue types and disease states within a single tissue section, allowing for a more precise deconvolution of genetic profiles and gene expression profiles than traditional bulk assays. While healthy tissue showed consistent gene expression across samples, OSCC micro-samples revealed large patient-to-patient variability in gene expression. KLK8 and KRTDAP only exhibited high expression in patient P3 (Fig-3a). KLK8 is implicated in malignant progression of OSCC [37], and KRTDAP strongly correlate with the differentiation and maintenance of stratified epithelium. Correspondingly, ‘keratin pearls’, structures indicating high-degree differentiation, were present in SRS images of P3 (Fig. S13) [38]. In addition to variability between patients, we found significant variability in gene expression between samples identified as cancer nests from the same patient. Specifically, in P4, the adjacent P4S1C and P4S2C micro-samples exhibited similar gene expression, however, P4S3C, which was recovered from a more distant location to previous two (Fig. S14), revealed a distinct gene expression profile including expression of CST1, which is related to promoted cell proliferation (Fig-3b). The intra-tumor molecular variation may indicate a branched evolution of tumorigenesis which could provide insight for tumor diagnosis and treatment.

Displacement and recombination of genes, especially oncogenes, has been recognized to drive neoplasia [39] and has become the focus of numerous cancer studies as potential therapeutic targets [40-42]. we then exploited *de novo* identification of gene fusion events through sequencing these samples. Early fusion events are usually sporadic and hence hard to identify with bulk sequencing. Furthermore, DNA sequencing of small samples with low input heavily relies on whole genome amplification, which is prone to chimera formation and causes false identification of fusion events. We used RNA-seq of micro-samples for *de novo* identification of gene fusion transcripts to investigate gene fusion events between patients and within tumor samples. Some fusion related genes were shared between micro-samples from separate patients, including KRT6A, which was shared between P1S2C and P3S5C, and FAM102A, which was shared between P1S3C, P3S5C, and P4S2C. Some samples, however, contained unique gene fusion patterns (Fig-3c, Sup. Excel 1.3-1.8). Most of these fusions might be passenger events that came along with cancer development and thus their actual consequences remain unknown. Among those, we found several sets that are consistent with previous observations, including one recorded in TCGA (MYH9 and KRT14, from P2), and two involving oncogenes AKT3 (fused with LRRC45, from P3) and MAFB (fused with SAC3D1, from P3) (Fig. S15, Supplementary Table 2, Sup. File1.1). Sanger sequencing on amplified fragments harboring joint junction of fused genes confirmed these fusions between MYH9 (5’ fusion partner: exon 20) and KRT14 (3’ fusion partner: exon 8), AKT3 (5’ fusion partner: UTR) and LRRC45 (3’ fusion partner: intron), implying the capability of SMD-seq in recurrent fusion event discovery. Fusion events also revealed substantial intra-sample heterogeneity (Fig. S16). For example, more fused genes were found in P4S3C, than in P4S1C and P4S2C which were dissected from distant regions of the tissue section, indicating spatially dependent genome instability during tumorigenesis. Moreover, we also observed the existence of an oncogene-involved fusion (RAB3D and MTMR14, verified by Sanger sequencing, Fig. S15, Sup. File1.1) in sample P4S2E (Fig. S17), which was identified as healthy epithelium by H&E and SRS analyses, again indicating the possibility that this microsample was progressing through the early stages of tumorigenesis.

In parallel to RNA-seq, DNA-seq was performed using half of the lysate from micro-samples, enabling the analysis of heterogeneity at the whole genome level. Copy number alteration (CNA) analysis demonstrated unique patterns of CNAs between cancer and normal samples, and between patients (Fig-4a, Fig. S18), indicating the high complexity in OSCC including significant genetic mosaicism and genetic heterogeneity. A few commonly shared large-size ploidy shifts, such as the losses of 3p and 8p and the gains of 3q and 8q, which are shared in other squamous cell carcinomas [45-46], were also observed in our OSCC samples. This analysis also revealed patient-specific CNAs, for example, chromosome 6 showed high instability in P2’s cancer sample but this instability was not present in others (Fig. S19-20). Unsupervised clustering of the samples based on CNAs further demonstrated the unique CNA patterns between patients (Fig-4b, Fig. S19) [43]. Micro-samples dissected from different locations within a patient sample displayed subtle discrepancies in copy number profiles (Fig-4b). The CNA pattern of chromosome 1 in P1S1C and P1S3C were different from that of P1S4C, with an obvious gain of 1q in the first two ROIs (Fig. S20).

**Fig-4.**
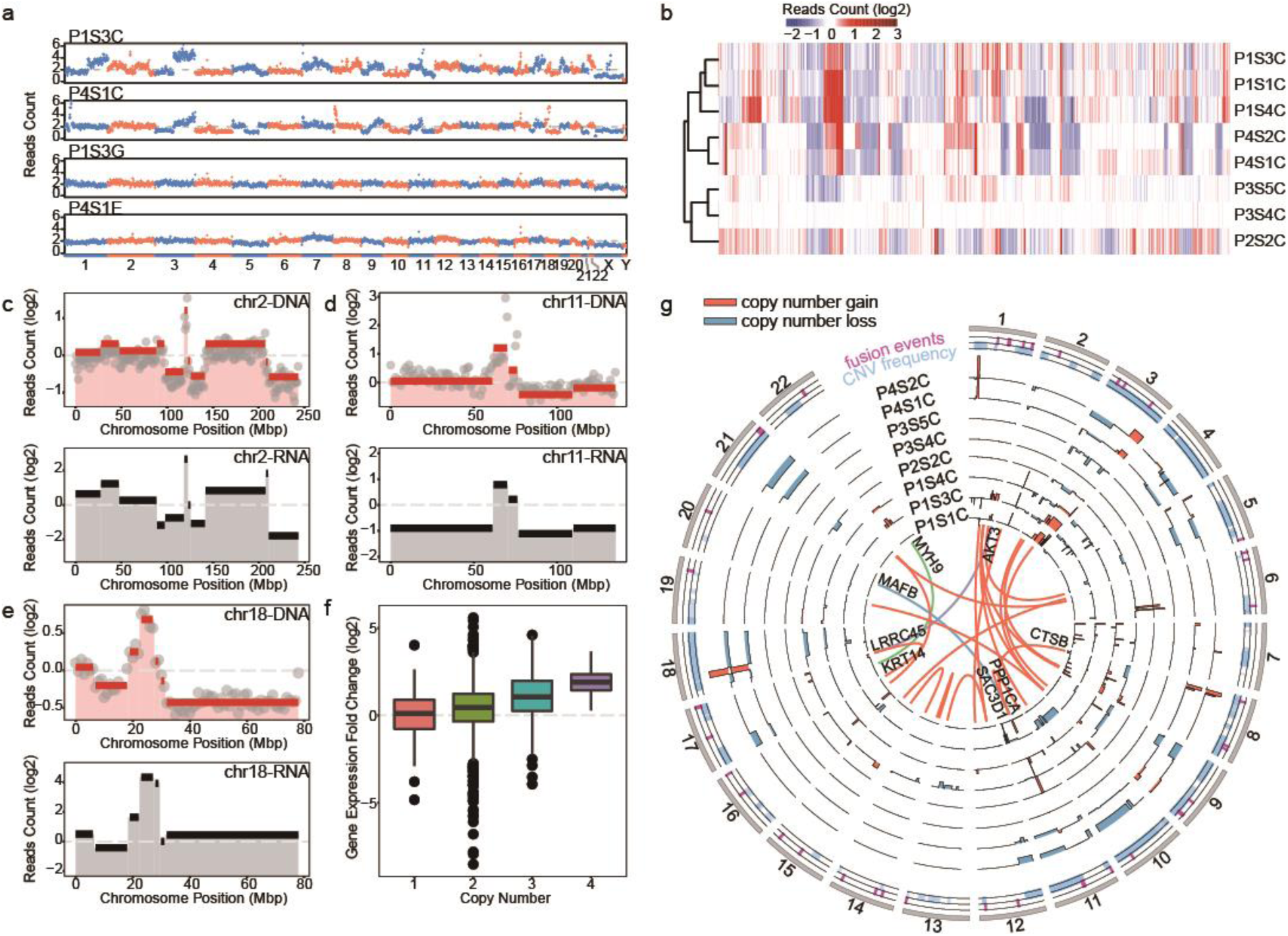
Genomic and transcriptomic analysis of laser dissected tissue samples. a) Raw reads count across the whole genome of two paired cancer-normal tissues dissected from the same slice. Chromosome numbers were labeled at the bottom. b) Unsupervised clustering of normalized reads counts of all the dissected cancer samples. c,d) Top panels demonstrated the normalized read count (grey dots) and copy numbers (red lines) identified by CBS algorithm in chromosome 2 and 11 of sample P1S1C and P1S3C, respectively. Mean expression levels within the same segment were shown in black lines in the bottom panel. e) Averaged read counts of all the cancer samples (top) and their corresponding gene expression values (bottom) of chromosome 18. Genome and transcriptome variations of other chromosomes and samples were shown in Fig. S16, 18-19. f) Averaged gene expression fold changes were computed per 1M bin across the whole genome and plotted against copy numbers. g) The distribution of CNV and fusion genes across the genome of all the cancer samples. Orange and blue bars indicated the copy number gains and loss, respectively. Orange lines in the inner circle indicated fusion genes with at least 10 span pair reads, and the green and purple lines represented oncogenes involved gene fusions. The position of fusion events (magenta) and CNVs (light blue) of all the cancer samples were shown in the 2 outmost circles.

As both genomes and transcriptomes were sequenced from each cancer nest, we were able to perform joint analysis of genomic and transcriptomic variation. The mean expression level of genes within each genomic segment [44] was compared with the copy number in the same regions (Fig-4c, d, e, Fig. S20,22), and summarized across the whole genome (Fig-4f, Fig. S23). The average gene expression levels showed positive correlation with the copy number within the same segment (Fig-4f). Among all the OSCC samples, 8 regions of recurrent copy number gain and 5 regions of recurrent copy number loss were identified (q < 0.25, Fig. S21, Sup. Excel 1.9-2.0).[45] Among these (copy number loss/gain or regions) 11q13.3, 8q24.3, 11p15.4 and 11q24.2 co-localized with differently expressed genes in cancer samples. GSTP1 located within the recurrent focal amplification of 11q13.3[46-47], which implied that the high expression level of GSTP1 may be the result of increased copy number. FAM83H was also co-localized with a focal amplification region, 8q24.3, and it specifically expressed at higher levels in patient P1. TP53AIP1 and PKP3 both expressed at a lower level in all the patients and located in the regions of recurrent copy number loss 11q24.2 and 11p15.4, respectively. The expression level of GSTP1, FAM83H, TP53AIP1 and PKP3 were all reported to be involved in the development of cancer or affecting patients’ survival rate [46-48]. We also analyzed the global correlation between gene fusion events and copy number variations and found a high-degree of overlap between them (Fig-4g). Fusion genes CTSB and PPP1CA co-localized with focal amplification regions 8p23.1 and 11q13.3, separately. CTSB was proved to be related to cancer progression and metastasis [49-50], and PPP1CA was reported to contribute to ras/p53-induced senescence [51]. Of the 24 pairs of detected fused genes, 17 pairs (∼71%) had at least one gene intersected with a focal copy number alteration (Sup. Excel 2.1). The parallel observation of genomic rearrangement and gene expression fold change may illustrate that the instability of the cancer genome led to gene fusion events which were more likely to occur within amplification and deletion regions (Fig-4g) [52].

## Discussion

SMD-seq takes advantage of SRS microscopy, which can capture chemical information without staining, and low-input sequencing to give high quality genome and gene expression data from well-defined histological regions. This technique enabled dissection of cancer heterogeneity across multiple measurement modalities, including morphology, genome alteration, gene expression, and gene fusion.

Obtaining quantitative information about genetic changes within cells in their native environment is challenging with traditional microscopy or genomic analysis alone. The major challenge hindering the progress of these studies is technical: although sequencing analyses, especially single cell sequencing enables the systematic identification of cell populations in a cancer tissue, most of such studies start with a dissociated cell suspension [53], which retains no spatial information from the sample of origin. On the other hand, current methods in surgical pathology lack the capability to efficiently isolate specific cell populations in complex tissues/tumors, which can confound molecular results. Spatially resolved transcriptomic profiling [54-55], including *in situ* hybridization (ISH) gene expression [56-57] and recently developed spatial barcoding methods [58-60]. Nevertheless, a main limitation of these methods is their lack of *in situ* pathological information, which cannot be overlooked for its importance of proper clinical applications.

The morphological information recovered by SMD-seq is reconstructed from chemical contrast, unlike gene expression in other spatial transcriptomic approaches [54-55]. Therefore, recovery of histological features does not rely on data reconstruction through gene expression patterns or spatial barcoding. Such independence between morphology and gene expression can lead to previously unobserved characteristics of tumorigenesis. The discordance between the morphology and genomic profile of micro-sample P4S2E served an example. The sample was identified as normal epithelium by SRS histology, H&E staining of an adjacent tissue-section. However, the gene expression profile of P4S2E clustered more closely with the cancer samples (Fig-2g), and both the genetic and transcriptomic features of this sample reflected a cancer-like pattern. Furthermore, genes previously reported to be significantly mutated in OSCC, such as FAT1, PPP2R1A, PTEN, HRAS, and CREBBP28, are also found in P4S2E (Fig. S24). This inconsistency between imaging and sequencing could imply that tumorigenesis signature reflected by gene expression profiles may arise before of morphological characteristics, such observation would be difficult to be captured and confirmed by histology reconstructed through expression patterns.

Although SMD-seq can preserve morphology and sequence information with high quality, there are still limitations. First, SRS imaging in SMD-seq can identify local tissue features, however, there is an intrinsic trade-off between the field of view and resolution of the SRS histology images. In this study, the lower numerical aperture of the objective lens compromised the sensitivity for minor change in subcellular structures, which hindered in depth image analysis. Increasing pixel density of image may reduce the noise level with higher spatial sampling rate (Fig. S25), but prolonged exposure times lead to a higher chance of sample ablation during imaging (Fig. S25) and bring higher risk of the RNA degradation during imaging. Another limitation of SMD-seq is the accessibility of SRS microscopy, which requires specialized light sources and sophisticated optical configurations making the technique difficult to implement in general biology and clinical labs. The integration of SRS microscopy into turn-key system should in principle facilitate adoption of SMD-seq in the clinics [61].

In summary, we have shown that SMD-seq can readily detect spatially dependent transcriptional variation and chromosomal alteration in unfixed tissue sections, and it can discover the subtle and rare genetic alteration events, such as gene fusions, copy number changes and differentially expressed transcriptome, with high sensitivity and accuracy, underlying different histological features. In this study, we performed bulk genomic analysis on the dissected micro-samples, allowing parallel DNA and RNA sequencing on the same ROI. This small-bulk pool-and-split approach masks the single-cell-level heterogeneity within the sample and blurs the relationship between the genome and transcriptome in single cells. SMD-seq could be extended to single cell analysis with careful dissociation of micro-samples. While they could limit the ability of multiomic characterization, such a development would provide higher resolution molecular profiling, including the analysis of epigenetic information. In combination with deep genome sequencing, the detection of SNVs can also be incorporated into current methods. SMD-seq offers a new direction for cancer research, with the integrated analysis of histology, transcriptome and genome, it has the potential to enable a more comprehensive understanding of the tumorigenesis process and diagnosis.

## Materials and Methods

### Pipeline of SMD-seq

#### 1, Preparation of tissue section

Tumor samples were collected from four patients who suffered various stages of OSCC at different ages and with different genders. All the biopsies were collected by protocols reviewed by and approved by the Ethics Committee of Peking University School and the Hospital of Stomatology (PKUSSIRB-201418116). Tumor samples diagnosed with oral squamous cell carcinoma were stored in DMEM supplemented with 10% (vol/vol) FBS at 4 °C immediately after operation. Samples were stored in ice bucket and transported to lab within 30 min. The samples were then frozen in a cryostat (CM1950, Leica, Germany) at −20 °C, and were sectioned to 30 μm thin sections. The sections were placed onto PEN membrane slides (RNAase-Free, Leica, Germany). All sections were kept on ice before imaging.

#### 2, SRS imaging and laser micro-dissection in situ

Each slide was surveyed with SRS microscopy at Raman peaks 2850 and 2950 cm^-1^ immediately after sectioning. The two Raman peaks represent CH2 symmetric vibration and CH3 vibration, respectively. To obtain a fast switch between the two peaks, the wavelength of optical parametric oscillator (OPO) was set at 813 nm (corresponding ∼ 2901 cm^-1^) and switching to 809.8 nm and 816.4 nm by tuning the lyot filter in the cavity. Each image was scanned with a pixel dwelling time of 4 μs, and a size of 640 × 640 pixels. To each image, a 2-frame Karlman filtering was applied. After above processing steps, the regions of cancerous cells were able to be identified by the experienced collaborating pathologist. A quick check at different focal planes was done before micro-dissection to make sure no big morphology change presenting within the 30 μm thickness. The post objective excitation power was 100 mW for pump, and 40 mW for Stokes when imaging and the pump (813 nm) power was increased to 180-190 mW to perform micro-dissection. To dissect an ROI, the laser repeatedly scanned 500 times along the border with a pixel dwelling time of 5 ms, hence effectively incised the polyethylene napthalate (PEN) membrane from the glass slide, facilitating the cutting of ROIs. The dissected sample was manually transferred from slide to tube loaded with lysis buffer by RNase-free syringe needle. For each tissue section, we selected at least one dissected region of tumor, and two dissected areas of similar size from normal tissues, one of them from the epithelium (the origin of this tumorigenesis) and the other from gland or muscle. To avoid the severe damage of RNA in the tissue sections, the whole procedure needs to be completed promptly.

#### 3, Transcriptome and genome sequencing

The dissected samples were put into lysis buffer [62] separately and immediately centrifuged at 13000 rpm for 30s. After lysis, each sample was equally divided into two aliquots for RNA-seq and genomic DNA sequencing, respectively. The protocol of RNA-seq was adapted from the pipeline of single-cell transcriptome analysis [62]. In brief, mRNA was reverse transcribed into first strand cDNA with polyT primer which has an anchor sequence. After other used primers were digested, polyA was added to the 3’ end of cDNA and second strand cDNA was formed and amplified with polyT primer with another anchor sequence by PCR. We employed degenerate oligonucleotide primed PCR (DOP-PCR) for amplifying the whole genome of each lysed tissue sample by the GenomePlex Single Cell Whole Genome Amplification Kit (WGA4-50RXN, Sigma-Aldrich, USA). For each sample, 50 ng of amplified genomic DNA and cDNA were used as the start amount of libraries preparation, separately. The pair-end sequencing libraries with ∼300 bp insert size were constructed following the instructions of NEB Next Ultra DNA Library Prep Kit for Illumina (E7370, New England Biolabs, USA). Illumina HiSeq 2500 systems were used for sequencing.

### Stimulated Raman scattering (SRS) microscope

The home-built SRS system used a pump laser integrated optical parametric oscillator (picoEmerald, APE, Germany). It provided two spatially and temporally overlapped pulse trains, with the synchronized repetition rate of 80 MHz. One beam, fixed at 1064 nm, was used as the Stokes light. The other beam, tunable from 780 to 990 nm, served as the pump light. The intensity of the Stokes beam was modulated at 20.2 MHz by a resonant electro-optical modulator (EOM). The overlapped lights were directed into an inverted multi-photon scanning microscope (FV1000, Olympus, Japan). The collinear laser beams were focused into the sample by a 20× objective (UPlanSAPO, NA 0.75, Olympus, Japan). Transmitted light was collected by a condenser (NA 0.9, Olympus, Japan). After filtering out the Stokes beam, the pump beam was directed onto a large area photo diode (FDS1010, Thorlabs, USA). The voltage from photo diode was sent into lock-in amplifier (HF2LI, Zurich Instruments, Switzerland) to extract the SRS signal. Image was reconstructed through software provided by manufacture (FV10ASW, Olympus, Japan).

### Image analysis

For two color SRS image, we applied a linear combination approach [63] on each field of view to convert the 2850 and 2950 cm^-1^ images into a reconstructed pseudo-color image (Fig-s1), in which we represented protein- and lipid-rich regions with blue and yellow, respectively. For texture analysis, all images were processed with the same procedure for HOG feature extraction. Dual color SRS images were resized to 80×80 pixels, and subjected to Matlab (Mathwork, USA) built-in function ‘extractHOGFeature’ to generate feature vectors. Hierarchical clustering was applied to extracted feature vector clustering in R, with built-in function ‘hclust’ (“ward.D” method). Image correlation matrix was generated in R.

### Sequencing data analysis

Reproducibility of SMD-seq of these samples were validated by checking the Spearman correlation coefficients (r) of expressed genes (average r = 0.7) and reads count (average r = 0.7) between biological replicates of the same patients. Adaptor contamination and low-quality reads (phred quality< 20) were discarded from the raw data. Only samples with coefficient of variation (CV) of reads count per 1M bin<0.25 (genomic DNA) and gene number more than 6000 (FPKM>0.1, RNA) were kept for analysis. For RNA-seq data, TopHat (v2.0.10) were used for sequencing alignment. Reference genome assembly hg19 and gene annotation files were downloaded from UCSC Genome Browser. FPKM values used for analyses were generated by Cufflinks (v2.1.1), and Cuffdiff (v2.2.1) was used for gene expression levels comparison. Significantly different expressed genes between muscle, gland, epithelium and cancer were selected under the criteria that p value < 0.05 and |log2 (fold change)| >1. Gene functional annotation was performed by The Database for Annotation, Visualization and Integrated Discovery (DAVID) v6.717 [64]. The purity of tumor samples was estimated by ESTIMATE [65] with gene expression data. They were submitted to ESTIMATE for calculation of each score, then pooled and compared with the database calculated from TCGA HNSCC samples (n=522) by the developer. Gene fusion analysis were carried out by FusionCatcher (v0.99.4a) [66] with four mapping tools (Bowtie, Bowtie2, BLAT, STAR). Matched normal samples were used for each patient to exclude the fusion genes that are also found in normal samples. Under following situations the fusion were discarded:(1) both fusion genes are mutual paralogs;(2) one or both of the fusion genes were pseudogene;(3) reported only by one mapping tool or reported by 2 mapping tools only once;(4) no known genes existed in between the fusion genes;(5) the distance between both genes were less than 100 kbp. Under this criteria, 24 fusion genes were discovered with more than 10 paired reads spanning two different genes sequences. The circular diagram of fusion gene was generated by CIRCOS (v0.67-7) [67]. RNA-seq data was used for variant calling by GATK (v3.4-0) according to GATK Best Practices recommendations [68]. We performed duplicate removal, SplitNCigarReads, base quality score recalibration before SNP calling, and filtered out SNPs by Fisher Strand values (FS>30.0), Qual by Depth values (QD<2.0) and sequencing depth passing the quality filter (DP<10). Annotation of SNPs was performed by SnpEff (v4.0) [69]. Significantly mutated genes in HNSCC (Head and Neck Squamous Cell Carcinoma) were inferred from COSMIC (Catalogue of Somatic Mutations in Cancer) and a comprehensive previous study [70]. Spearman correlation coefficient was computed between tissue samples by function ‘cor’ in R. The unsupervised hierarchical clustering was performed by the function ‘pheatmap’ of package ‘pheatmap’ in R. and the method of measuring the distance in clustering columns was ‘manhattan’. 3D PCA plot was generated by R package ‘scatterplot3d’.

The sequencing depth of DNA-seq was ∼0.1×. Genomic DNA sequencing reads were mapped to reference genome by bowtie2 (v2.2.3) [71]. After duplication removal of mappable reads, the counts of aligned reads were calculated in each 1M bin along the genome (Figure 5A, C, D, E). For each bin, the read count of each tumor sample was normalized by sequencing depth and the median read count of all normal tissue samples, and the generated copy number went through segmentation by Circular Binary Segmentation [72] (the significance level was set as 0.05). The mean gene expression values of cancer samples within each segment were also calculated and normalized by mean expression values of normal samples which had corresponding qualified (CV<0.25) gDNA reads (Figure 5C, D, E). Function “pheatmap” in R was adopted for CNVs clustering (Figure 5B). GISTIC 2.030 was adopted to analyze the significantly reoccurring focal alterations for the gDNA segmented data.

## Supplementary Materials

### Materials and Methods

Fig. S1. Reconstruction process of dual-color SRS histology image.

Fig. S2. Dual-color SRS images of Warthin’s tumor and mucoepidermoid carcinoma.

Fig. S3. SRS images of epithelium and cancer for unsupervised hierarchical clustering.

Fig.S4. Correlation matrix of samples in 16 cancer samples and 16 epithelium samples.

Fig. S5. Images of cancer infiltrating muscle of P4.

Fig. S6. Localization of SRS subimage in corresponding HE.

Fig. S7 Characterization of micro-dissection of micro tissues.

Fig. S8 RNA preservation comparison between H&E stained and unstained sections.

Fig. S9. Spearman correlation coefficients calculated between SMD-seq samples.

Fig. S10. Unsupervised hierarchical clustering and gene annotation of enriched genes in different tissues.

Fig. S11. Gene expression levels of AKR1B10, FTH1, FTL between cancer (C, orange) and epithelium samples (E, cyan) from different patients.

Fig. S12. Immunofluorescence images to show the protein expression level of GSTP1.

Fig. S13. Keratin pearls in cancer nest of P3.

Fig. S14. 3-dimensional location of cancer nests of P4.

Fig. S15. The validation of gene fusion events by Sanger sequencing.

Fig. S16. Fusion events of intra-tumor ROIs.

Fig. S17. Gene fusion events in sample P4S2E.

Fig. S18. Genomic sequencing coverage across the whole genome.

Fig. S19. Comparison of unsupervised clustering of normalized reads count between Ginkgo and our methods.

Fig. S20. Copy number variation of autosomes from all cancer samples.

Fig. S21. Significant focal copy number alterations of all the cancer samples analyzed by GISTIC 2.0.

Fig. S22. Mean gene expression fold change of autosomes.

Fig. S23. Copy number variation and gene expression fold change of the same sectioned slice.

Fig. S24. Significantly mutated genes in OSCC discovered by previous study and COSMIC.

Fig. S25. Effect of image size and sample damage caused by laser.

Table S1. Patients information and corresponding dissected tissues.

Table S2. Summary of RNA-seq datasets.

Table S3. Summary of genomic DNA sequencing datasets.

Data file S1. Summary of genomic sequencing and RNA sequencing data analysis

## Acknowledgments

We thank BIOPIC and Peking University Sequencing Center for the experimental help and sequencing service.

## Author contributions

Y. H., T. L. conceived the project; Y. H., T. L., T. C., C. C., and J. Z. designed all experiments; T. C. performed SRS imaging, *in situ* microdissection and image analysis; C. C. performed molecular biology experiments and bioinformatics analysis. J. Z. performed histology identification, H&E staining and immunofluorescent staining. T. C., C. C., J. Z., Y. H. and A.S. analyzed the data and wrote the manuscript.

## Data and materials availability

The sequence of SMD-seq samples have been deposited in the NCBI Sequence Read Archive (http://www.ncbi.nlm.nih.gov/Traces/sra/) under accession number SRP075236.

