## supplemental information 1 for "Histologically resolved spatial multi-omics of human oral squamous cell carcinoma": Manuscript-SI-biorxiv.pdf

**List of Supplementary Materials**

Materials and Methods

Fig. S1. Reconstruction process of dual-color SRS histology image.

Fig. S2. Dual-color SRS images of Warthin's tumor and mucoepidermoid carcinoma.

Fig. S3. SRS images of epithelium and cancer for unsupervised hierarchical clustering.

Fig.S4. Correlation matrix of samples in 16 cancer samples and 16 epithelium samples.

Fig. S5. Images of cancer infiltrating muscle of P4.

Fig. S6. Localization of SRS subimage in corresponding HE.

Fig. S7 Characterization of micro-dissection of micro tissues.

Fig. S8 RNA preservation comparison between H&E stained and unstained sections.

Fig. S9. Spearman correlation coefficients calculated between SMD-seq samples.

Fig. S10. Unsupervised hierarchical clustering and gene annotation of enriched genes in different tissues.

Fig. S11. Gene expression levels of AKR1B10, FTH1, FTL between cancer (C, orange) and epithelium samples (E, cyan) from different patients.

Fig. S12. Immunofluorescence images to show the protein expression level of GSTP1.

Fig. S13. Keratin pearls in cancer nest of P3.

Fig. S14. 3-dimensional location of cancer nests of P4.

Fig. S15. The validation of gene fusion events by Sanger sequencing.

Fig. S16. Fusion events of intra-tumor ROIs.

Fig. S17. Gene fusion events in sample P4S2E.

Fig. S18. Genomic sequencing coverage across the whole genome.

Fig. S19. Comparison of unsupervised clustering of normalized reads count between Ginkgo and our methods.

Fig. S20. Copy number variation of autosomes from all cancer samples.

Fig. S21. Significant focal copy number alterations of all the cancer samples analyzed by GISTIC 2.0.

Fig. S22. Mean gene expression fold change of autosomes.

Fig. S23. Copy number variation and gene expression fold change of the same sectioned slice.

Fig. S24. Significantly mutated genes in OSCC discovered by previous study and COSMIC.

Fig. S25. Effect of image size and sample damage caused by laser.

Table S1. Patients information and corresponding dissected tissues.

Table S2. Summary of RNA-seq datasets.

Table S3. Summary of genomic DNA sequencing datasets.

Data file S1. Summary of genomic sequencing and RNA sequencing data analysis

### Materials and methods

*Large area imaging*

Image with large field of view was accomplished by multipoint time lapse function integrated in microscope software (FV10-ASW, Olympus, Japan). The outline determined by scanning and registering images along the boundary of ROI (region of interest). The software planned a scanning path according to the selected outline. Then, the stage automatically moved the sample to follow the planned path and acquired the image sequence with their spatial information. Each frame took ~21 s to obtain an image of 1024×1024 pixels. For the large field-of-view label-free histological image, two scanning sequences were required for lipid and protein channel.

*Immunofluorescence imaging*

Fresh tissues were embedded with OCT tissue-freezing medium, Frozen sections (5 μm thick) were blocked with a solution containing 2.5% bovine serum albumin for 30 minutes at 37 °C. Subsequently, sections were incubated with GSTP1 (3F2) Mouse mAB (1:200, Cell Signaling Technology) at 4 °C overnight. The secondary antibody, goat anti-mouse IgG-Cy3 (Invitrogen), was applied. Nuclei were stained with DAPI. Slides were washed on glycerol (1:1) for examination. Immunofluorescent signals were viewed using a confocal laser scanning fluorescent microscope (ZEISS LSM 5 EXCITER laser scanning microscope, Carl Zeiss MicroImaging, Oberkochen, Germany).

*PCR, cloning and transfection*

The primers used for amplifying fusion junction fragments from extracted cDNA were listed in Supplementary Table 2. PCR was performed with following cycles: 94°C 3 mins, 35 cycles of 94 °C 30 s, 60 °C 30 s, and 72 °C 30 s, followed by 72 °C 5 mins. The PCR products were run on 2% agarose gel and gel-purification with Zymoclean Gel DNA Recovery Kit (D4008, Zymo, USA). Recovered DNA were cloned into pMD18-T vector (D101A, Takara, Japan), and transfected to DH5α competent cells (CD201, Transgen, China). After incubation overnight in 37 °C, single clones were picked, and subjected to PCR (primers were listed in the following Table) verification and Sanger sequencing (Supplementary Figure 9).

Table. Primers used for amplifying the fusion sequence.

| Primer Name | Target Fusion Gene | Primer Sequences (5'-3') |
|---|---|---|
| MK-F | MYH9-KRT14 | CTGCCTACCTGAAGCTGCG |
| MK-R | | AGGACCTGCTCGTGGGTG |
| AL-F | AKT3-LRRC45 | AGCAGCAGCAGAGAATCCAA |
| AL-R | | CTCTCCCTGTCCAGCAGC |
| RM-F | RAB3D-MTMR14 | GGCGCCTCTTTCTCAGGTCC |
| RM-R | | ATCCCAGCAAAGATGGGGTGG |

*Comparison between H&E stained and unstained cryosections*.

Eight 30-μm-thick OSCC cryosections were prepared successively after snap-frozen (fig. S6). One of them was for H&E staining, and the other 7 were kept unstained. 12 ROIs of 3 tissue types (cancer, epithelium, and muscle) were micro-dissected from the H&E stained section and subjected to cDNA extraction, with 4 ROIs of each tissue type. For unstained sections, 4 ROIs of cancer area were micro-dissected from each section every 20 minutes (T0-T6), and followed by cDNA extraction. Fragments of 3 housekeeping genes (GAPDH, β-Actin and PPIA) were amplified for qPCR (*Takara* Bio SYBR Premix Ex Taq, Clonetech, Japan). The experiment was performed twice.

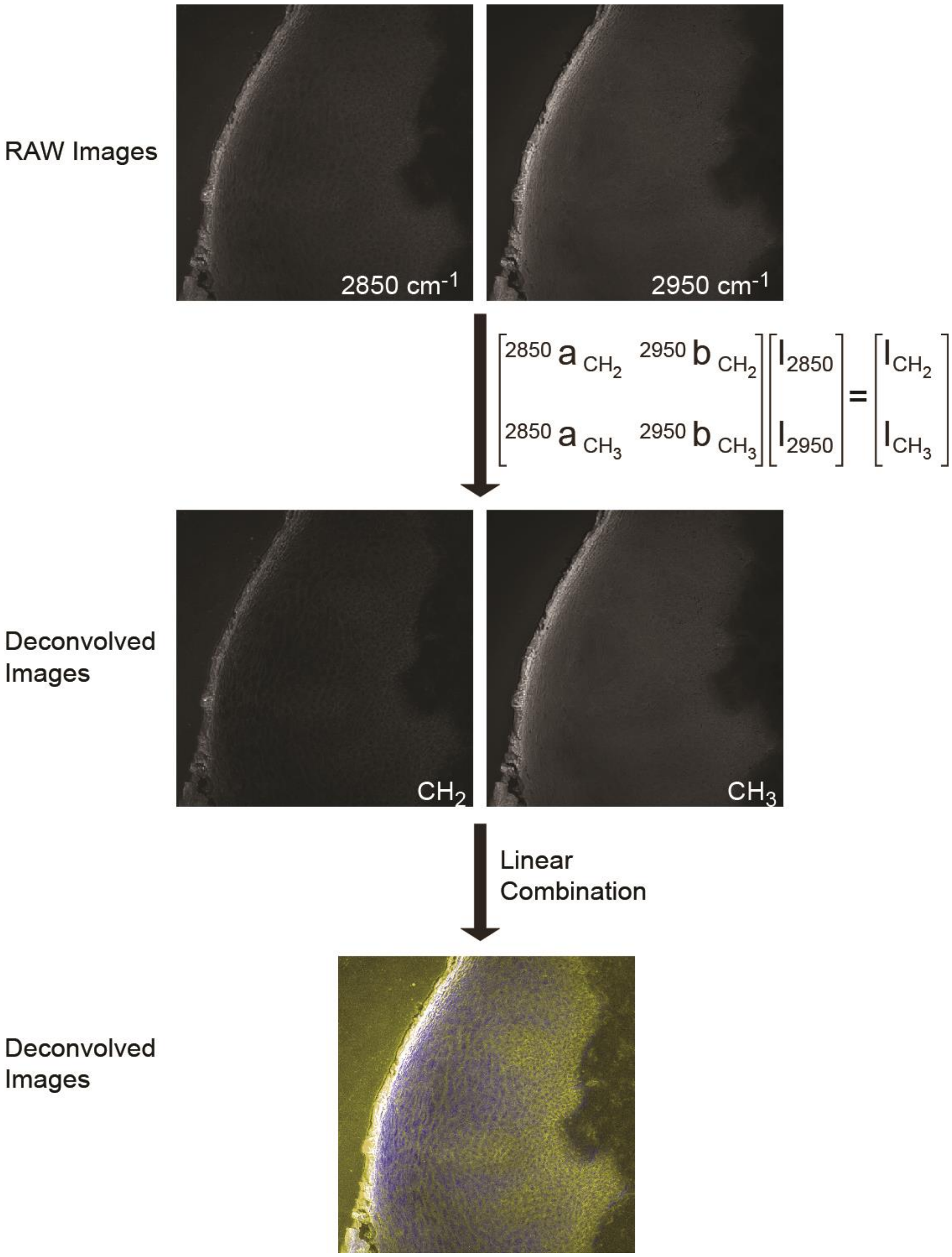


$$\begin{bmatrix} {}^{2850}a_{CH_2} & {}^{2950}b_{CH_2} \\ {}^{2850}a_{CH_3} & {}^{2950}b_{CH_3} \end{bmatrix} \begin{bmatrix} I_{2850} \\ I_{2950} \end{bmatrix} = \begin{bmatrix} I_{CH_2} \\ I_{CH_3} \end{bmatrix}$$

**Fig. S1** Pseudo color SRS histological image reconstruction process.

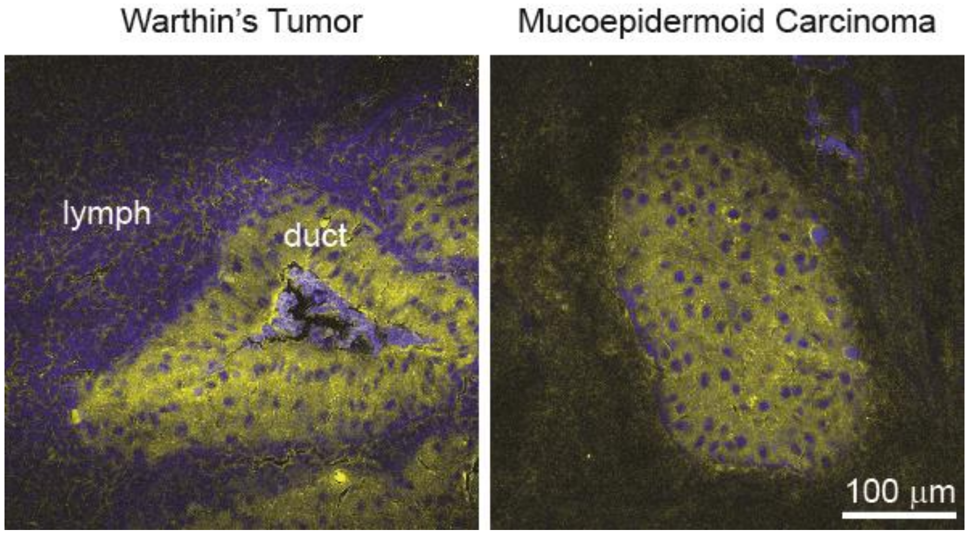


**Fig. S2** Dual-color SRS images of Watson tumor and mucoepidermoid carcinoma.

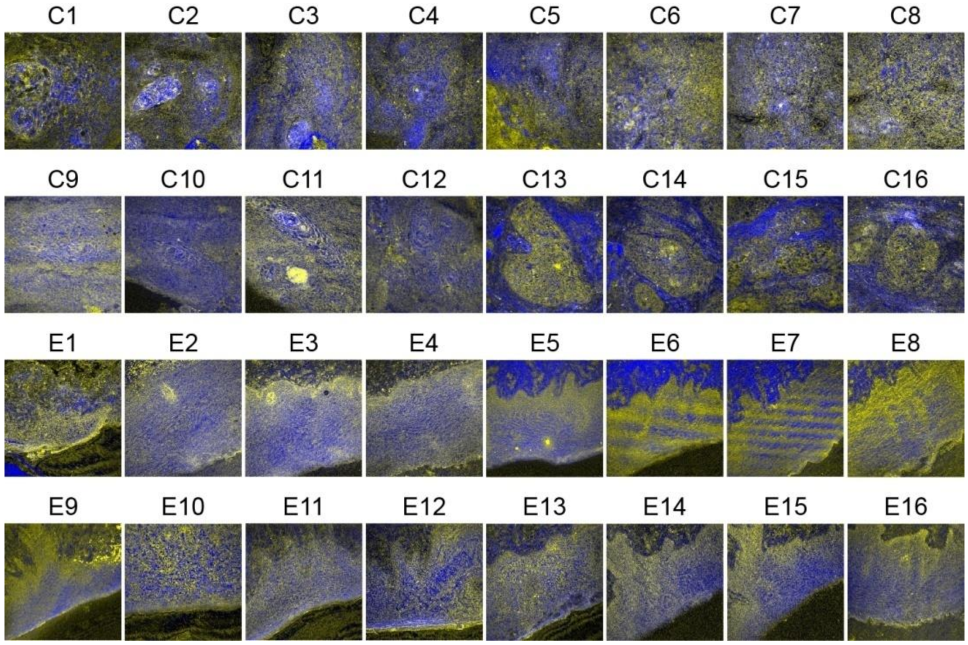


**Fig. S3** SRS images for unsupervised hierarchical clustering. 16 images of cancer samples (C#) and 16 images of epithelium samples (E#) are shown. Epithelium samples were visually inspected and manually adjusted with orientation.

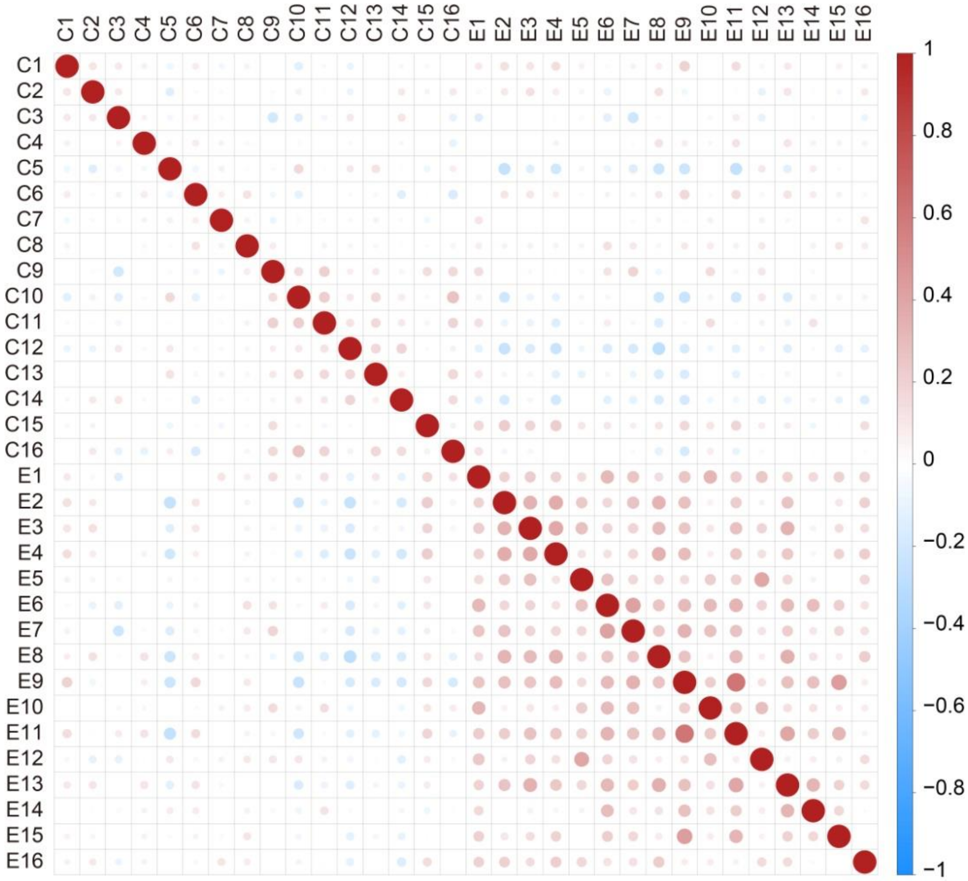


**Fig. S4** Correlation matrix of samples in 16 cancer samples and 16 epithelium samples (Supplementary Figure 5). Each dot represents the correlation coefficient of HOG features between two samples.

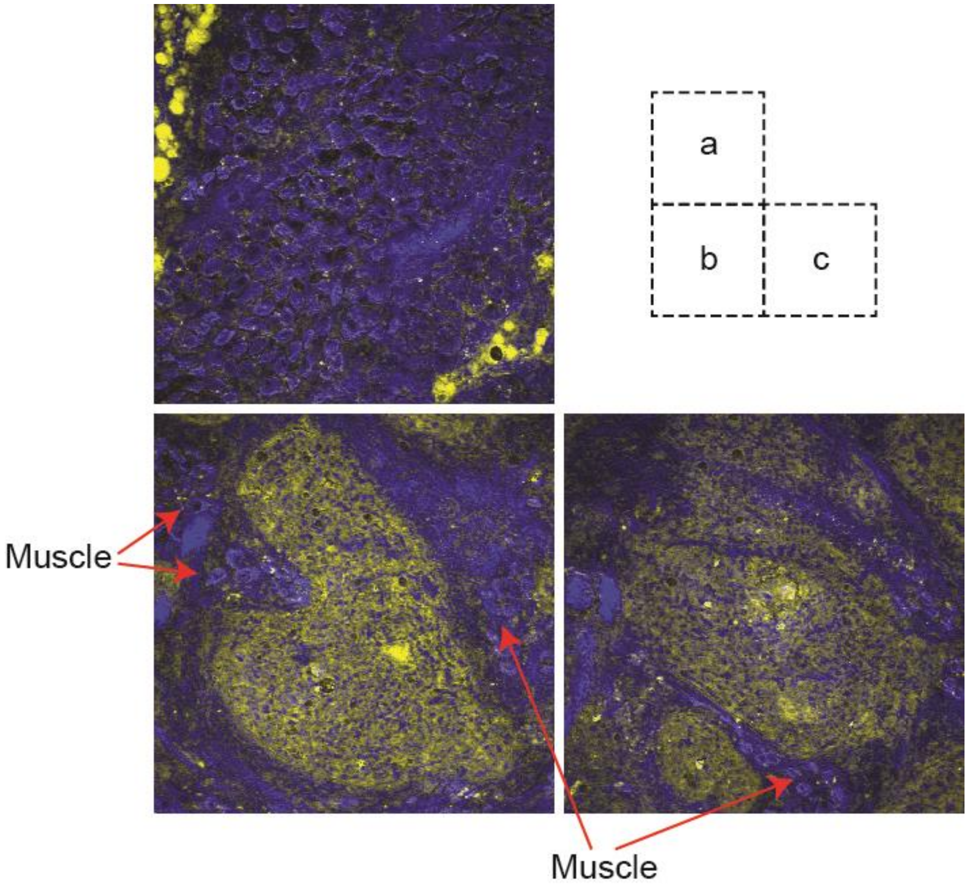


**Fig. S5** The muscle near cancer nests of P4. a) typical SRS image of muscle tissue, presenting high protein content. b,c) two cancer nests of P4, presenting muscle, which has been infiltrated by cancer, as marked by red arrows.

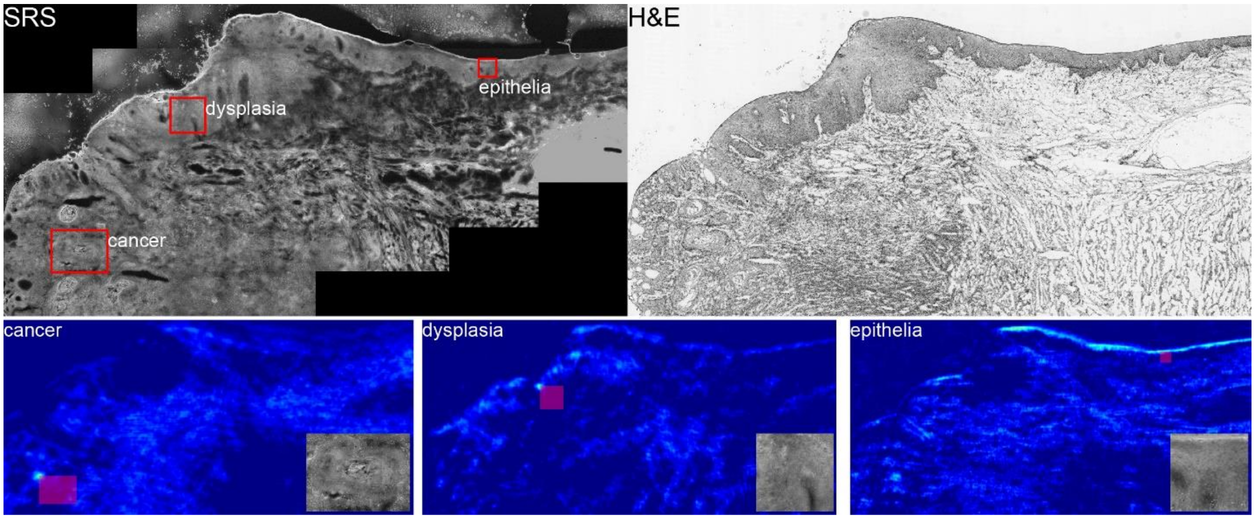


**Fig. S6** Localization of SRS subimage in corresponding HE. The first row presented stitched SRS image and corresponding H&E, both in gray scale. Three areas were selected (white dashed boxes) for localization in H&E image. The second row showed the correlation map between HOG features of selected SRS subimage and H&E stitched image. The highlighted spot in correlation map is the localized reference point of the subimage. It is notable that the epithelia presented a line of high correlation, indicating much higher structure conservation of the normal tissue.

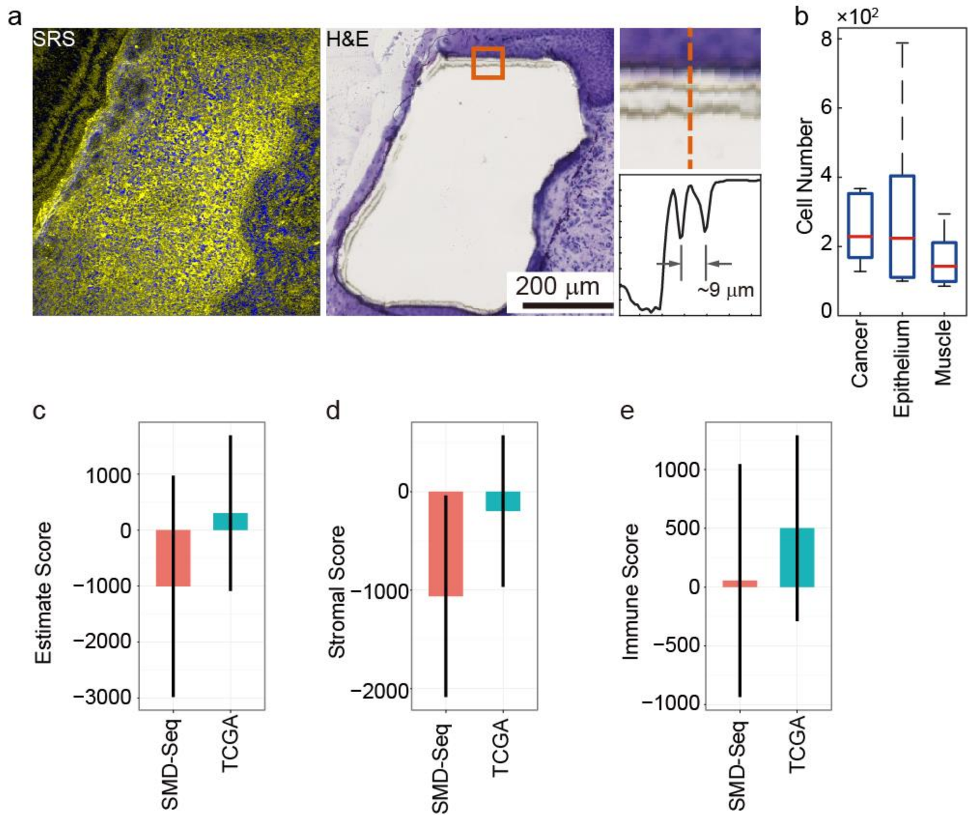


**Fig. S7** Characterization of micro-dissection of micro tissues. a) SRS image and H&E staining after microdissection of the same epithelium microsample. The magnified micrograph showed the width of incision line, which is about 9 µm. b) the statistics of cell number in collected microtissues. c-e) SMD-seq micro-tissues overall sample purity, stromal and immune cell contamination compared with TCGA samples.

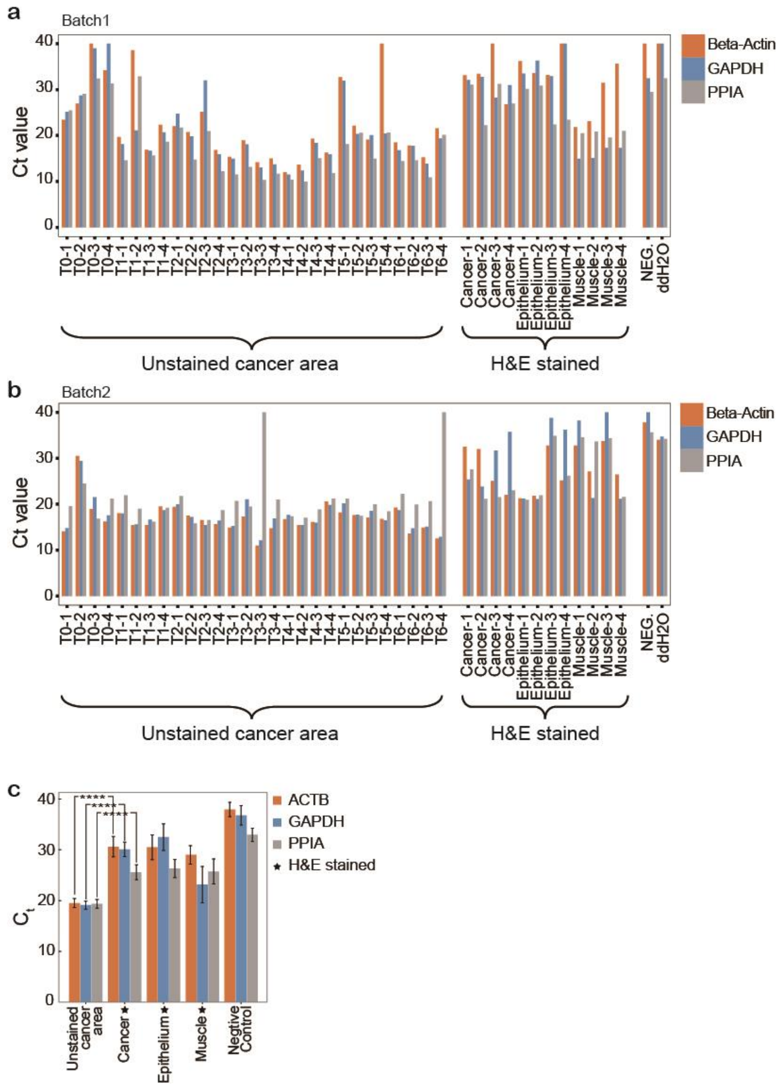


**Fig. S8** RNA preservation comparison between H&E stained and unstained sections. a-b) qPCR Ct values of housekeeping genes, including beta-actin, GAPDH, and PPIA, in two batches of experiments. Each batch contained 7 different time points for unstained cancer sample. c) statistics showed the different between RNA preservation different before and after staining.

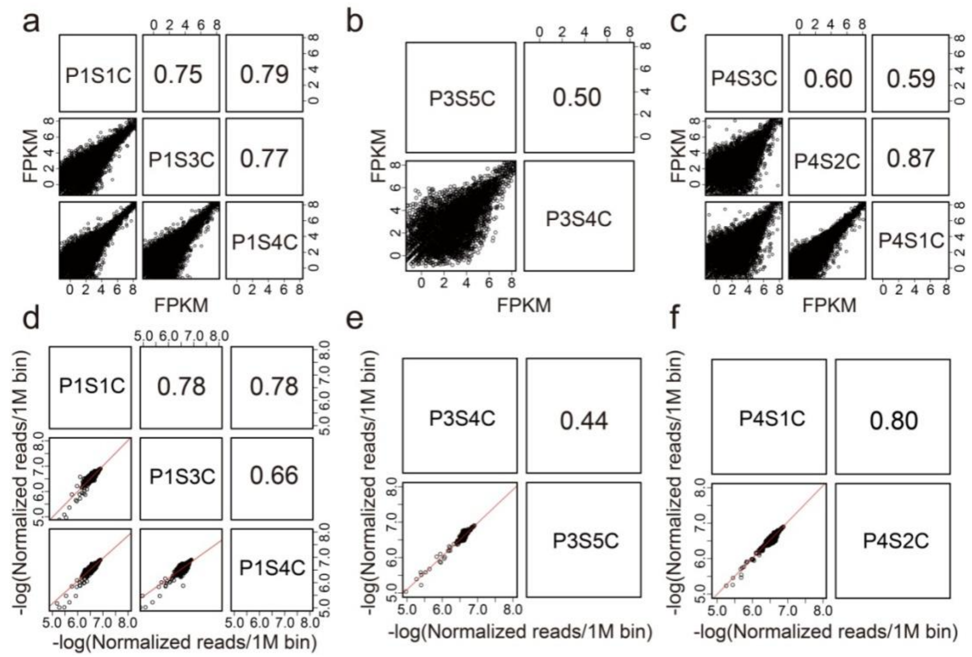


**Fig. S9** Spearman correlation coefficients calculated between SMD-Seq samples. Each dot in (a-c) represents the FPKM value (FPKM > 0.01) of one gene by RNA-Seq. Reads mapped in each 1M bin were normalized against total sequencing depth in DNA-Seq as plotted in (d-f).

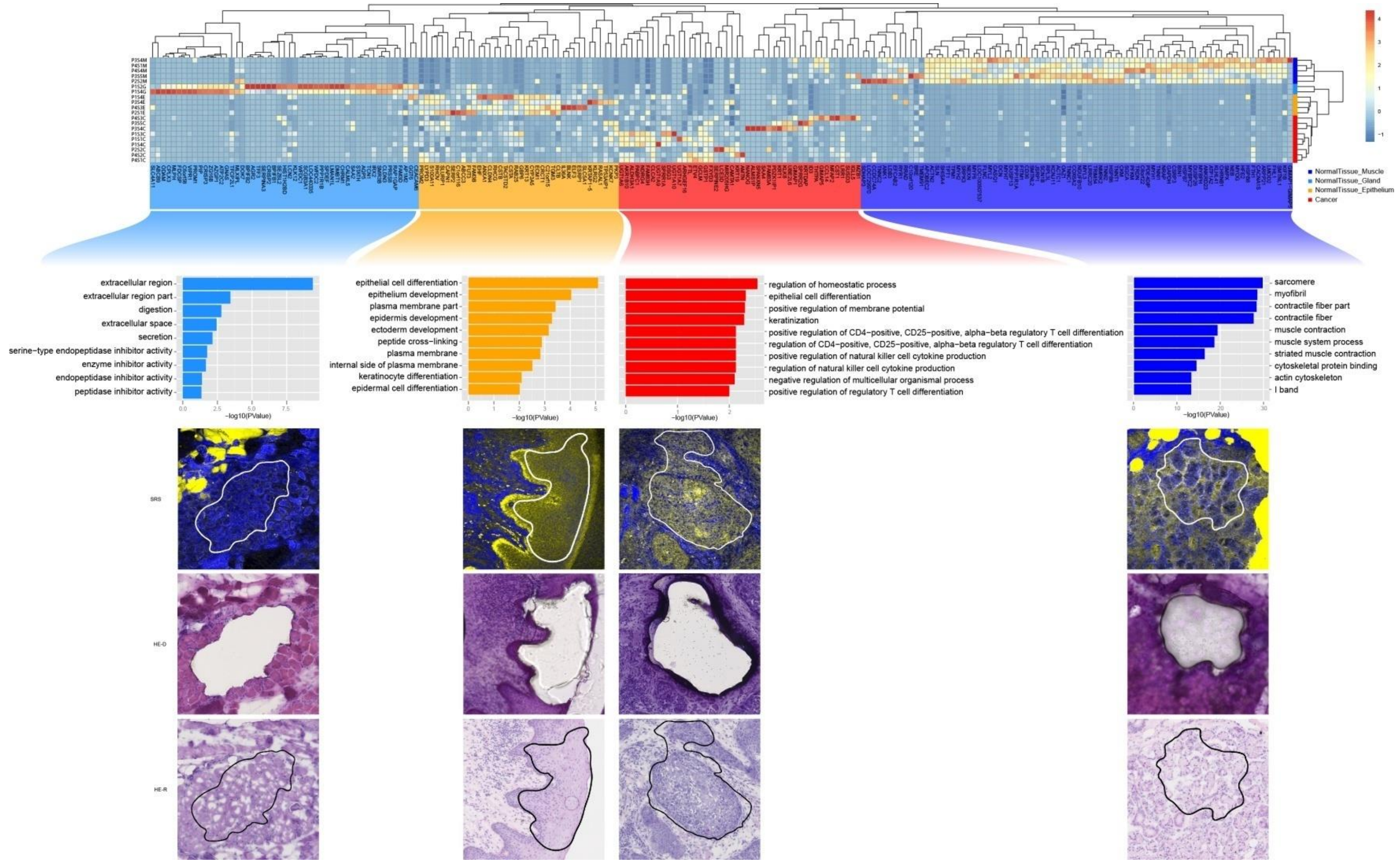


**Fig. S10** Unsupervised hierarchical clustering and gene annotation of enriched genes in different tissues. Unsupervised hierarchical clustering of differently expressed genes in epithelium and cancer. Top 10 GO terms with P-value < 0.05 are shown. GO terms were ranked by logarithm P-value. Typical images, including SRS and H&E staining, of each tissue type were shown.

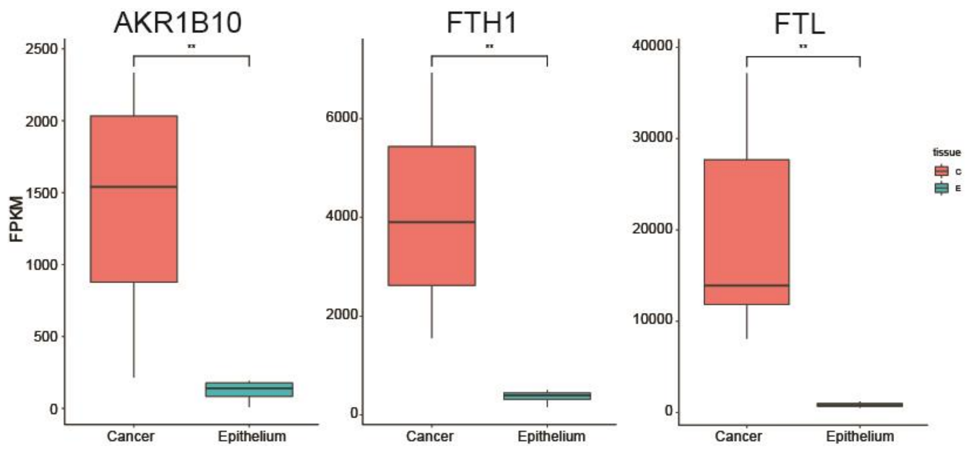


**Fig. S11** Gene expression levels of AKR1B10, FTH1, FTL between cancer (C, orange) and epithelium samples (E, cyan) from different patients.

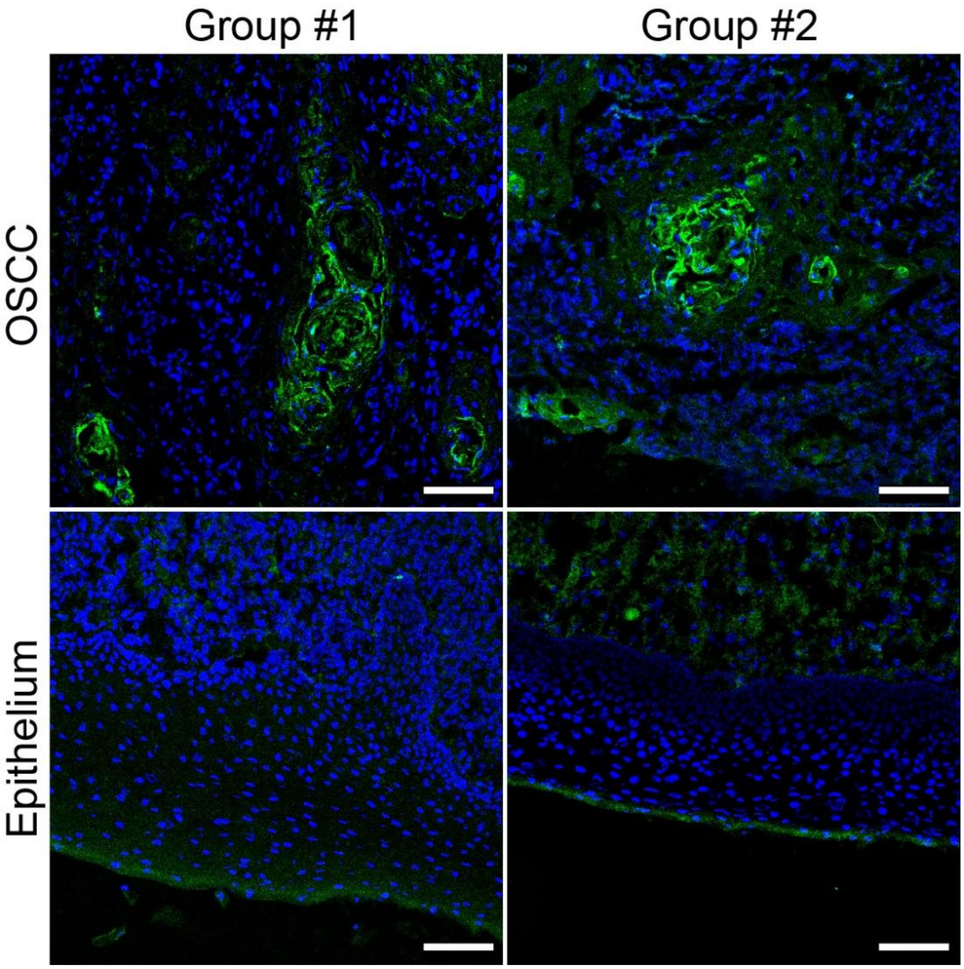


**Fig. S12** Immunofluorescence images to show the protein expression level of GSTP1. Green channel is fluorophore linked to protein antibody, representing GSTP1 positive region. Blue channel is DAPI, representing the neucleic. Group1 and group2 were replicates. Scale bar is 100 μm.

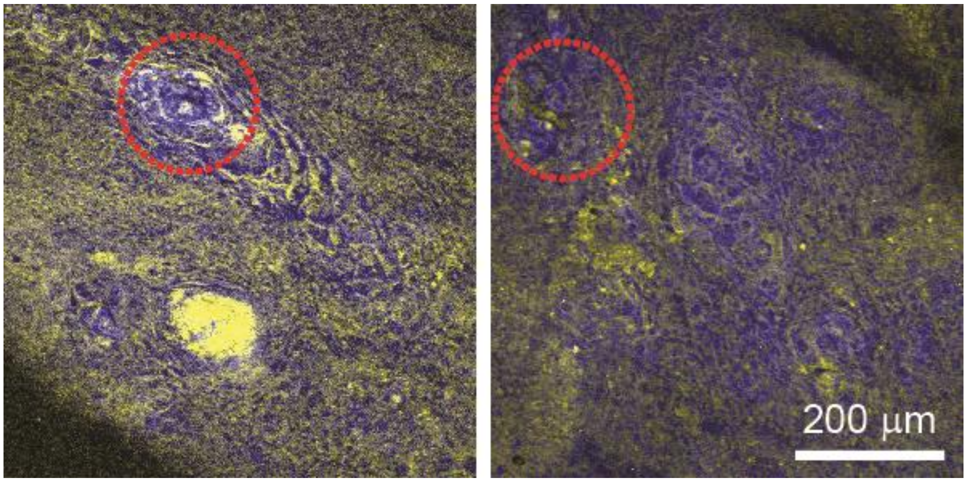


**Fig. S13** Keratin pearls in cancer nests of P3, marked by red circles.

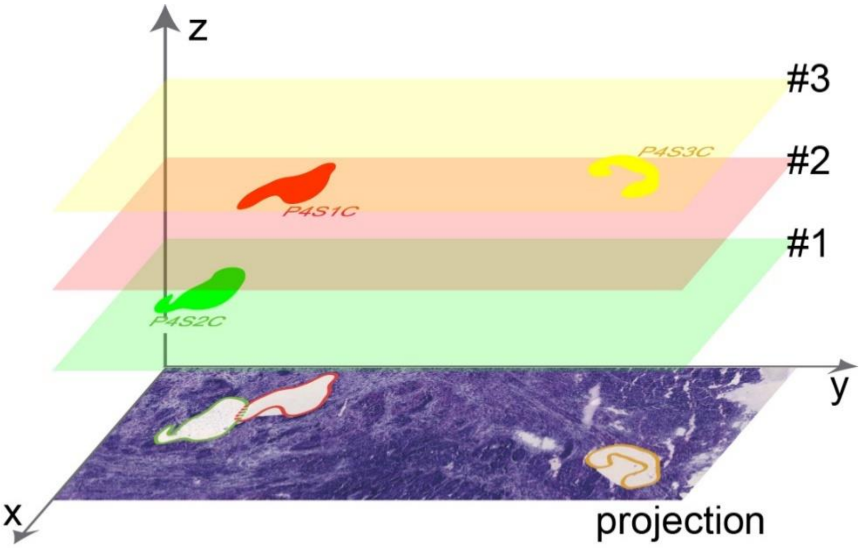


**Fig. S14** 3-dimensional locations of cancer nests of P4.

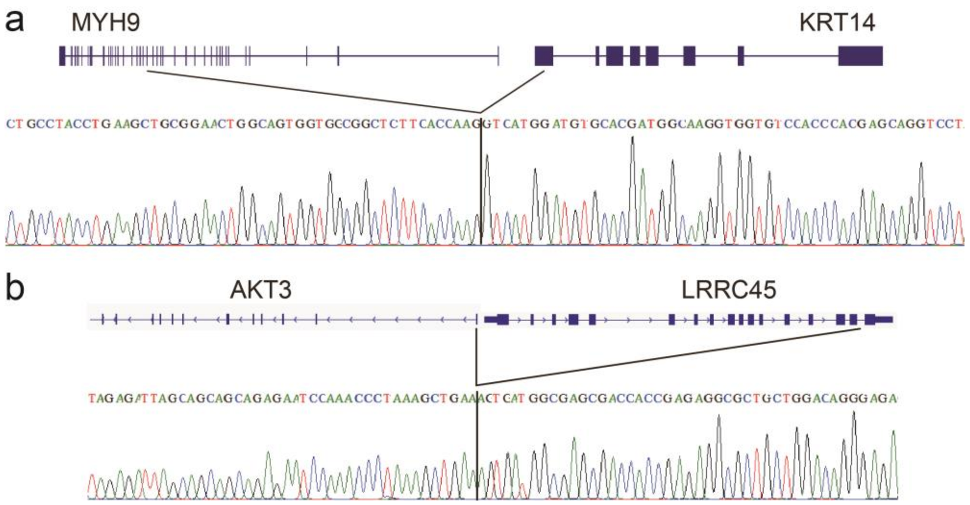


**Fig. S15** The validation of gene fusion events by Sanger sequencing. (a) Fusion of MYH9 and KRT14. Black lines connect the fusion parts of two genes. (b) Fusion of AKT3 and LRRC45.

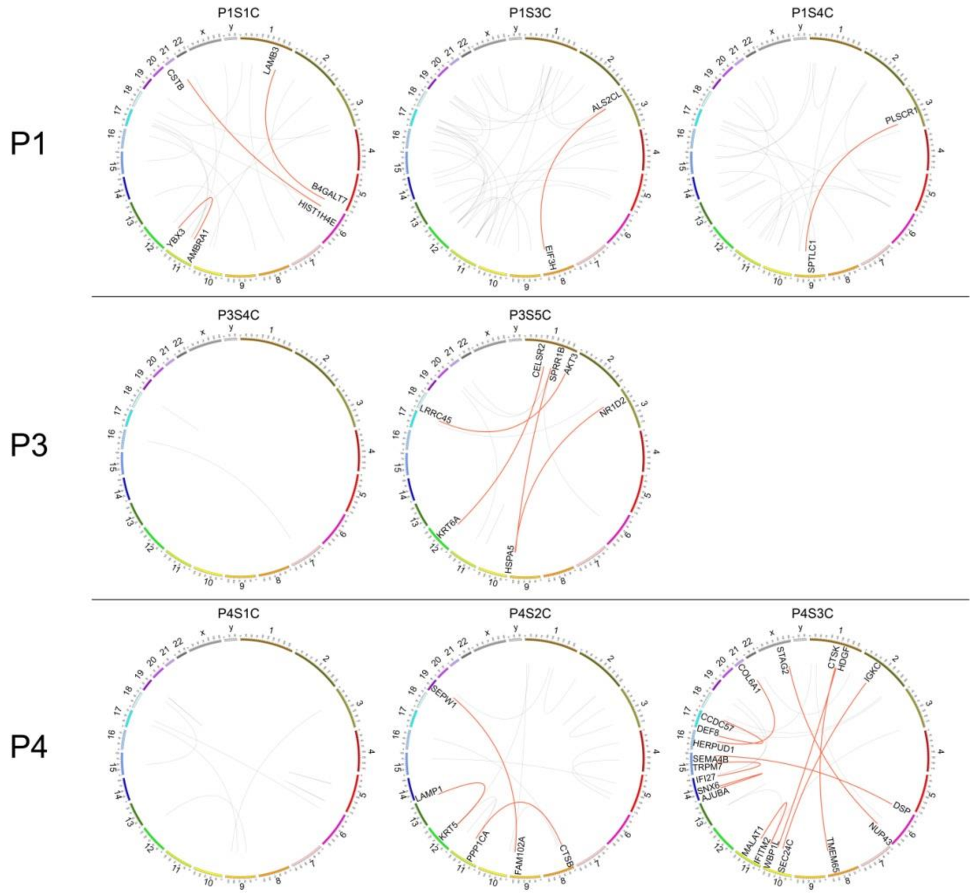


**Fig. S16** Fusion events of intra-tumor ROIs. Orange lines indicated fusion genes with at least 10 span pair reads, grey lines represented the other fusion genes.

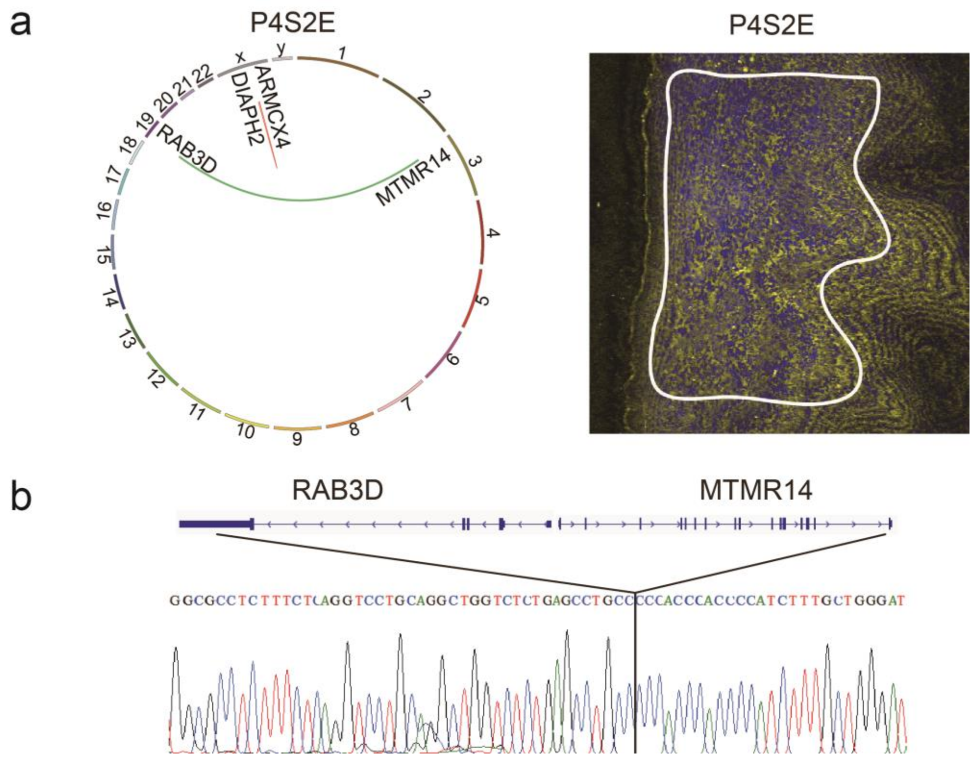


**Fig. S17** (a) Diagram of gene fusion events of P4S2E and its SRS image. The ribbons represent the fusion gene pairs. Fusion gene with more than 10 mapped span pair reads is shown in red, and oncogene involved fusion is colored in green. (b) Fusion of RAB3D and MTMR14.

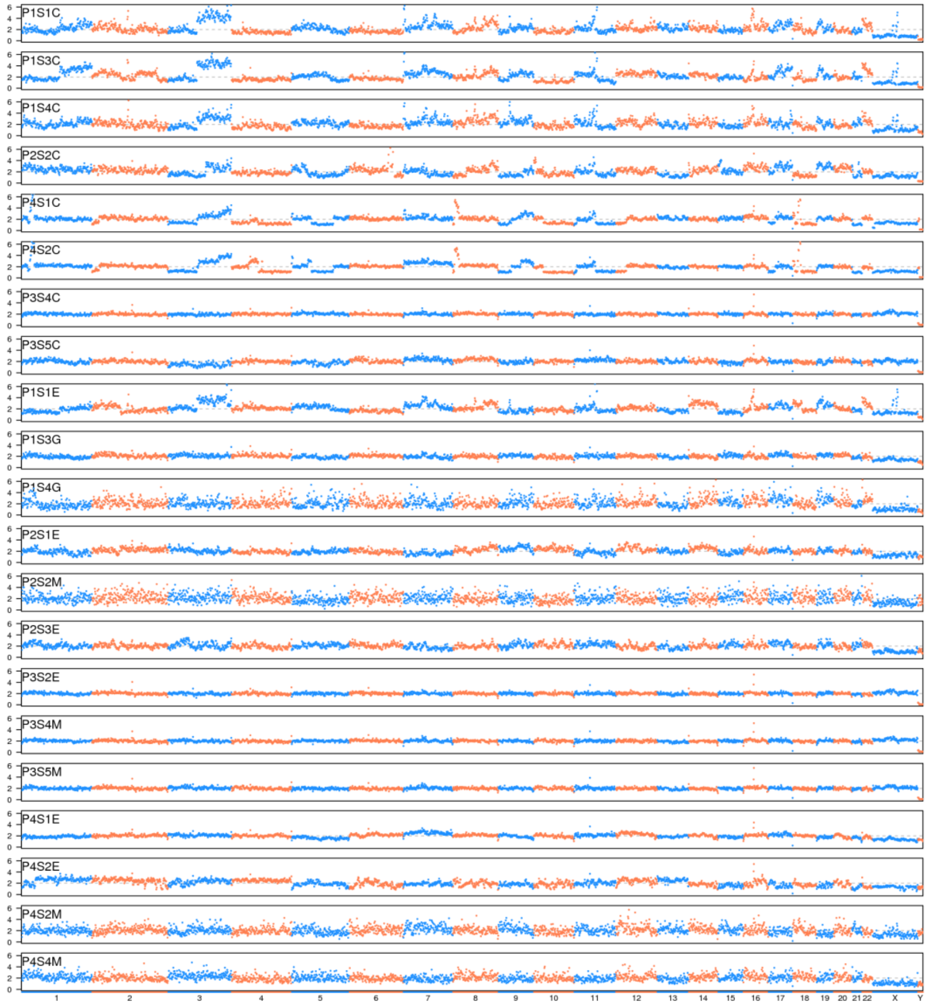


**Fig. S18** sequencing coverage across the whole genome, numbers at the bottom represent the chromosome numbers.

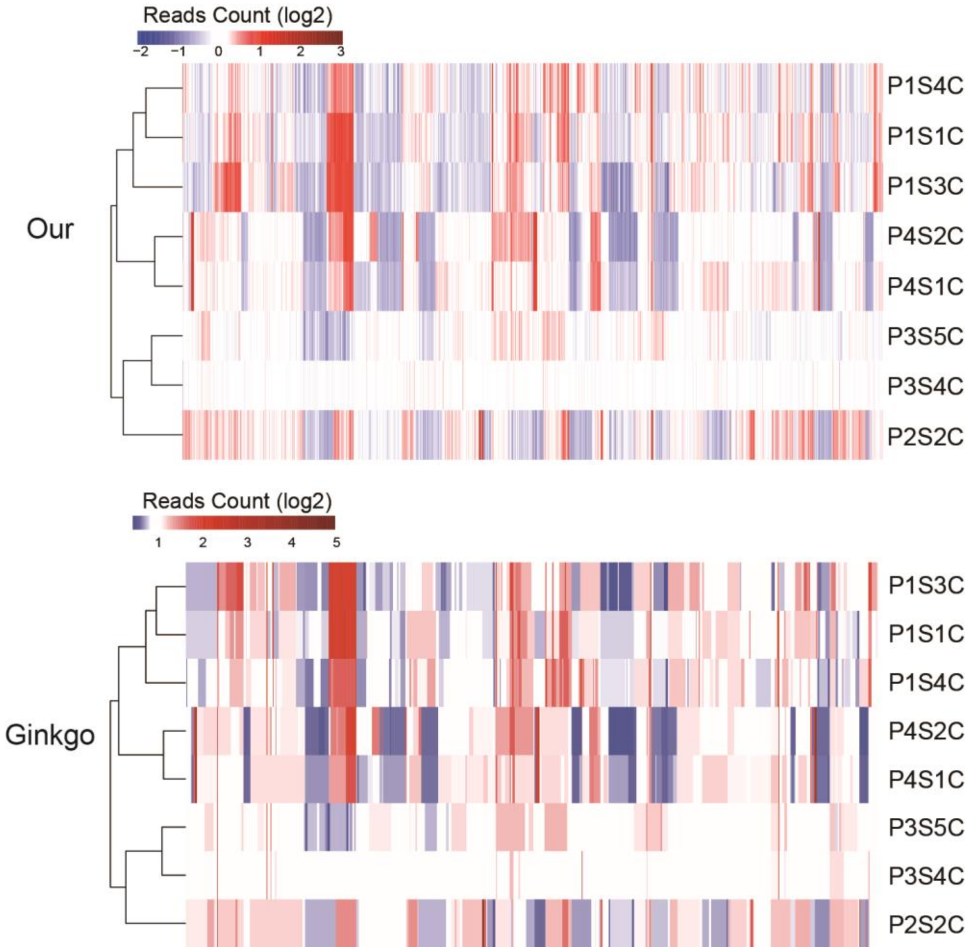


**Fig. S19** Comparison of unsupervised clustering of normalized reads count between Ginkgo and our methods. Ginkgo Both of them shows the similar clustering results in which samples from the same patient were clustered.

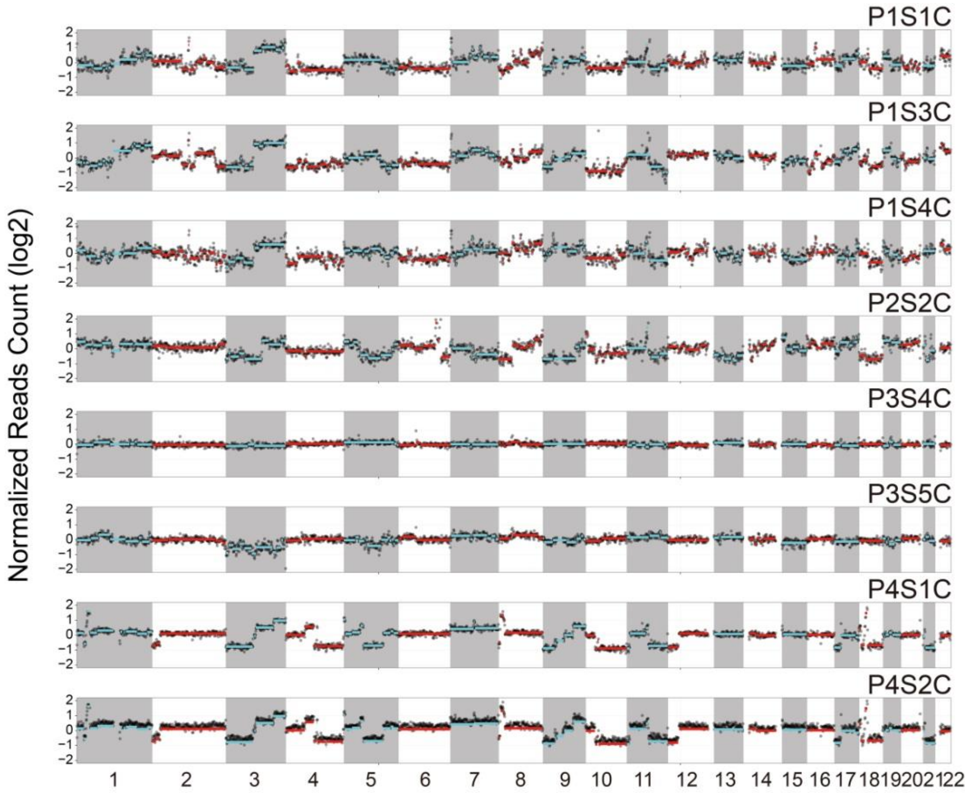


**Fig. S20** Copy number variation of autosomes from all cancer samples. Grey dots represent the normalized logarithm fold change, red and cyan lines demonstrate the segments along each chromosome calculated by CBS algorithm.

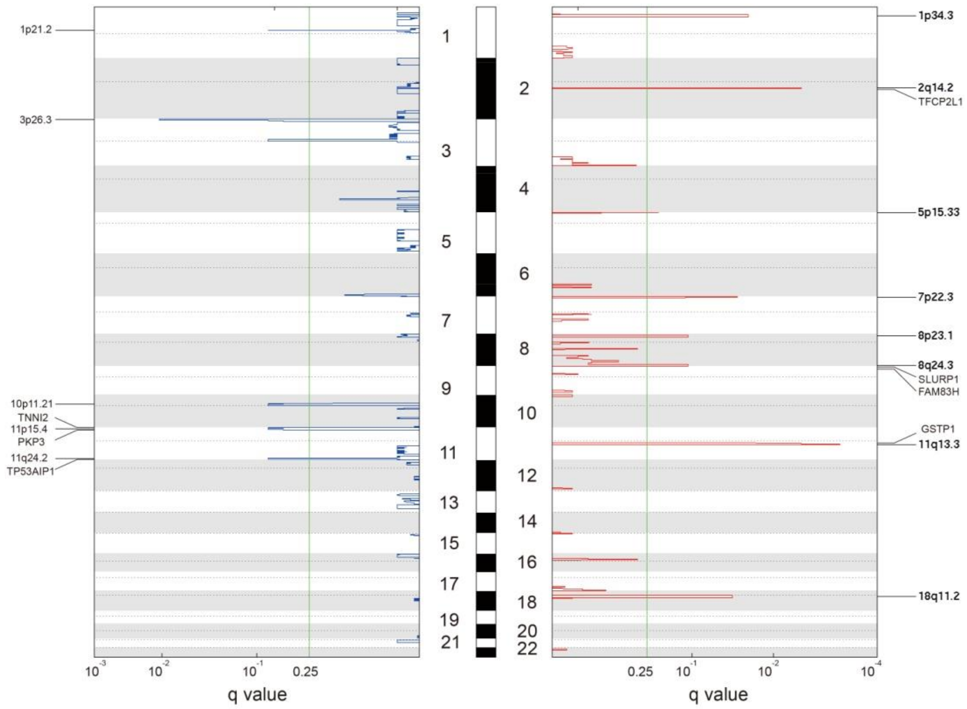


**Fig. S21** Significant focal copy number alterations of all the cancer samples analyzed by GISTIC 2.0. Red and blue lines represented amplification and deletions peak regions, separately. Amplification or deletion regions with < 0.25 q value were annotated with possible driver genes which were also identified in RNA-Seq as differently expressed genes.

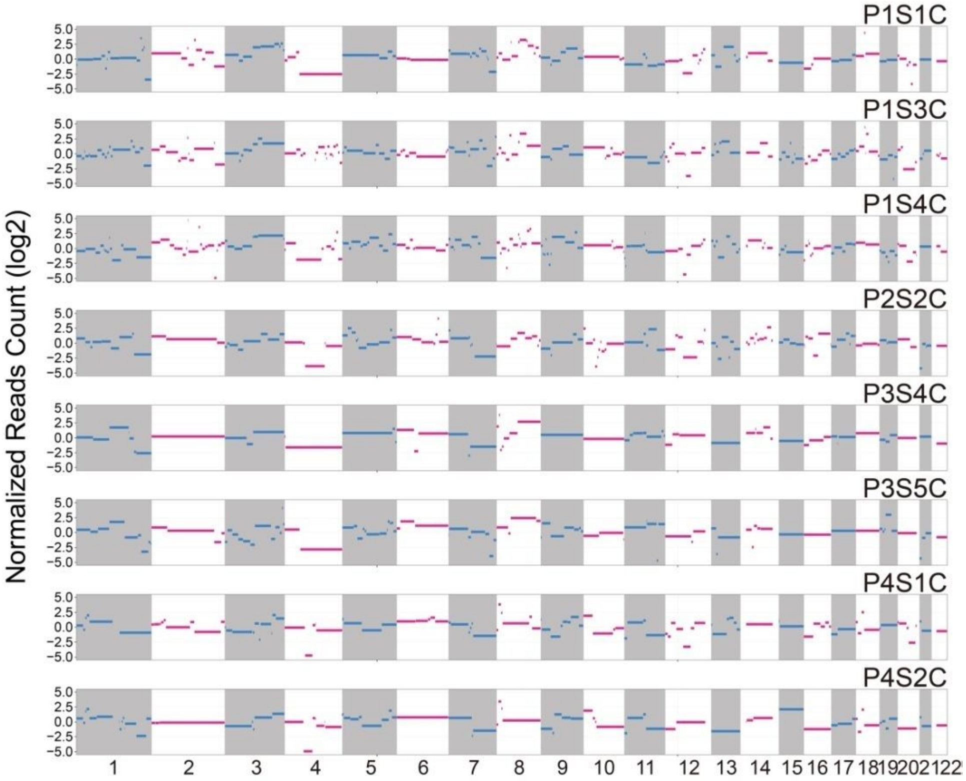


**Fig. S22** Mean gene expression fold change of autosomes. Magenta and cyan lines were mean gene expression value within each segment, which were calculated by CBS algorithm with normalized reads number per 1M bin.

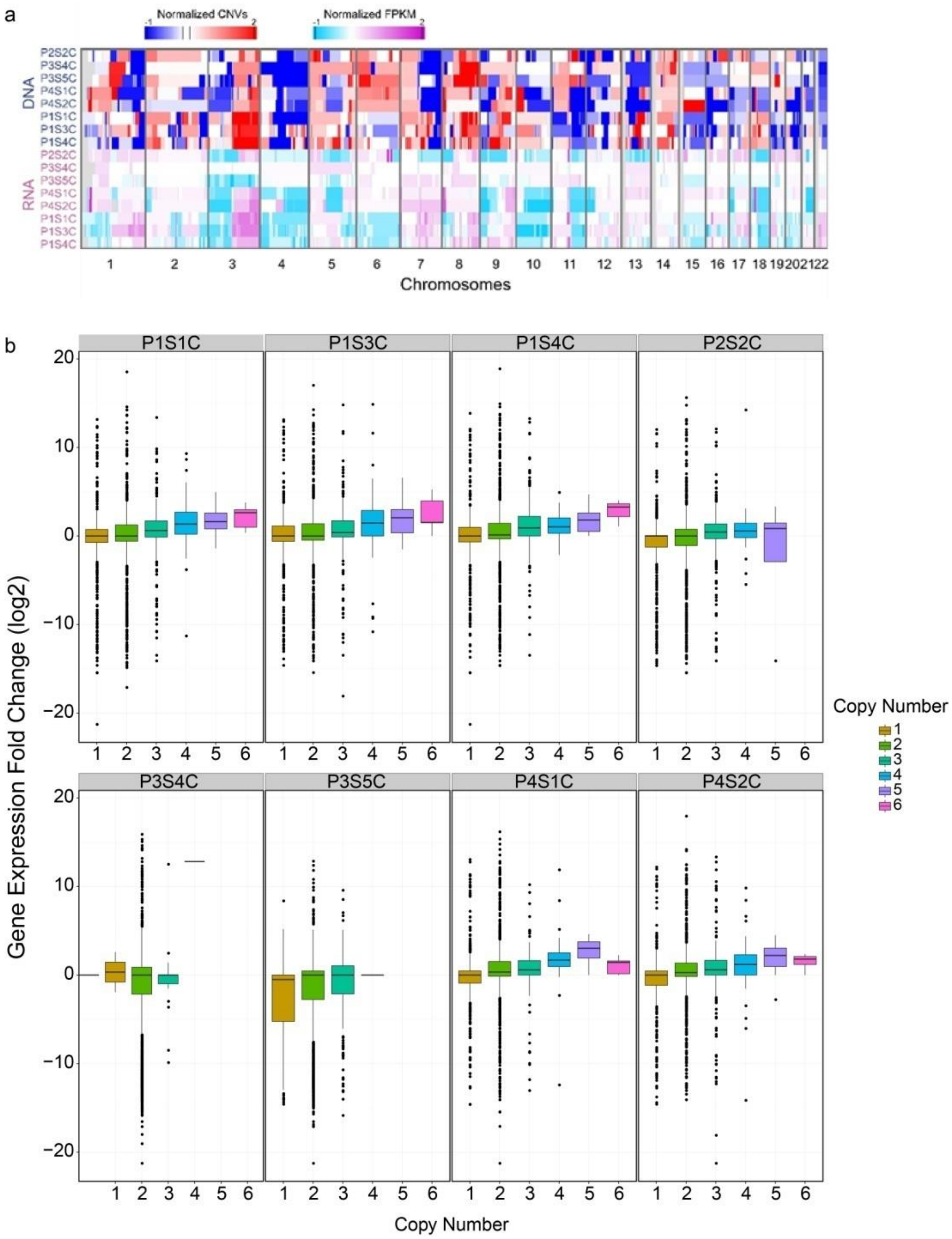


**Fig. S23** Copy number variation and gene expression fold change of the same sectioned slice. (a)number variation and gene expression fold change of the same sections. (b)Normalized gene expression level of cancer samples. Mean gene expression levels were calculated in each 1M bin along the genome. The copy number of each bin and its corresponding gene expression fold change are plotted.

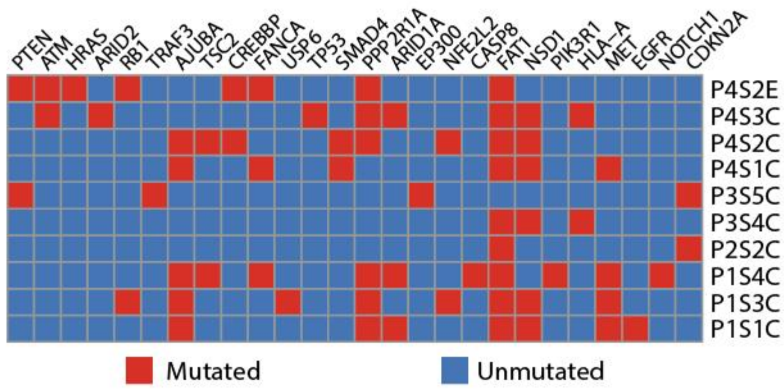


**Fig. S24** Significantly mutated genes in OSCC discovered by previous study and COSMIC. Red indicated the gene mutated in corresponding samples, blue demonstrated there was no SNP found.

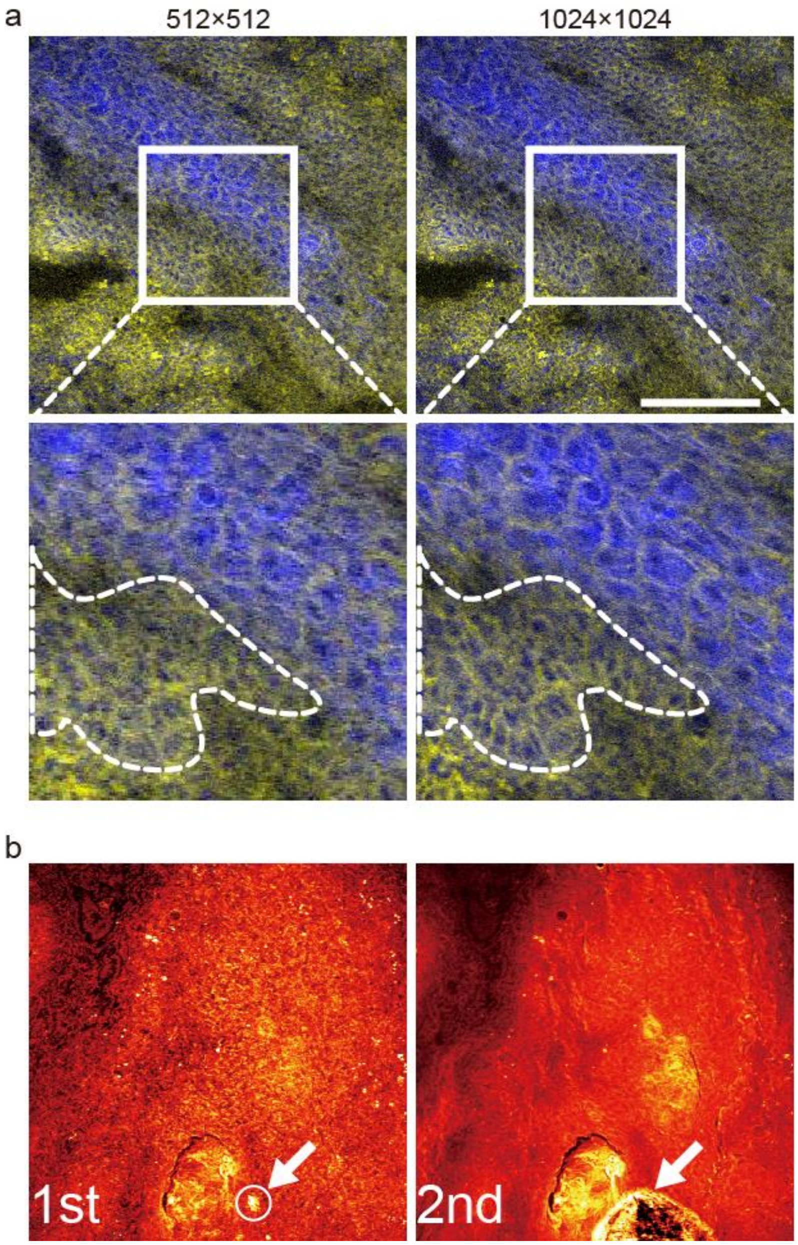


**Fig. S25** Effect of image size and sample damage caused by laser. (a) The same field of view imaged with different image sizes of 512×512 and 1024×1024. Cropped region covering the area was shown to see the difference. The cellular profile (closed white dash line) was clearer in the 1024×1024 image crop. Scale bar is 200 μm. (b) laser induced sample damage. The same field of view was imaged twice. In the first image, a bright spot could be seen (white circle, pointed out by arrow). It was the primary damage position. In the second image, the primary spot turned into a spreading down damage zone (pointed by white arrow).

**Supplementary Table 1** Patients information and corresponding dissected tissues. The first column shows 2-color SRS histological image, second column demonstrates the same tissue staining by H&E after laser dissection, and the last one represents a 5 μm thick HE stained tissue which is next to the section for SRS imaging.

| Patients | Dissection Date | Gender | Normal Epithelium | Normal Gland/Muscle | Cancer | Tumor Size | Location | Diagnosis |
|---|---|---|---|---|---|---|---|---|
| P1 | 2014-10-16 | Male | 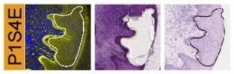  | 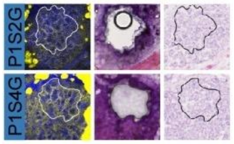  | 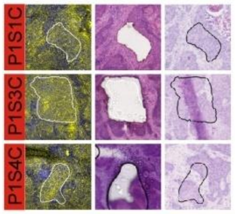  | 2 cm | The floor of mouth | OSCC; well-moderately differented |
| P2 | 2014-12-22 | Male | 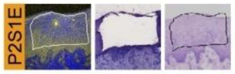  | 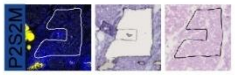  | 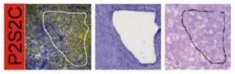  | 1.5 cm | Left tongue | OSCC; well-moderately differented |
| P3 | 2014-12-23 | Female | 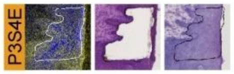  | 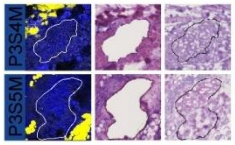  | 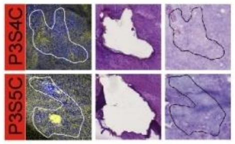  | 1.5 cm | Left tongue | OSCC; well-moderately differented |
| P4 | 2014-12-30 | Male | 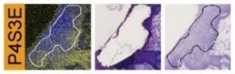  | 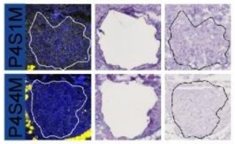  | 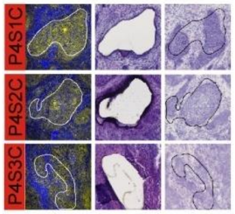  | 4 cm | Right tongue | OSCC; moderately differented |

**Supplementary Table 2** Summary of RNA-Seq datasets

| Sample | Raw reads | Q20 Reads | Clean reads | Mapped Reads | Mapping ratio | Gene Number (FPKM≥0.1) | Note |
|---|---|---|---|---|---|---|---|
| P1S1C | 12,069,236 | 8,467,776 | 7,272,185 | 6,822,412 | 93.82% | 10,542 | |
| P1S3C | 12,630,764 | 10,284,498 | 9,348,026 | 8,505,179 | 90.98% | 11,605 | |
| P1S4C | 19,732,916 | 14,904,428 | 12,891,237 | 11,595,447 | 89.95% | 11,932 | |
| P1S1E | 17,974,844 | 9,629,392 | 1,973,553 | 1,510,299 | 76.53% | 1,677 | Discard |
| P1S2G | 11,587,380 | 7,312,150 | 5,770,257 | 4,839,789 | 83.87% | 8,548 | |
| P1S3G | 13,630,336 | 8,988,472 | 6,569,778 | 5,962,025 | 90.75% | 4,310 | Discard |
| P1S4E | 15,154,164 | 8,008,098 | 3,373,990 | 3,006,803 | 89.12% | 7,982 | |
| P1S4G | 14,046,476 | 10,185,428 | 7,732,608 | 6,905,709 | 89.31% | 7,492 | |
| P5C1 | 19,800,126 | 15,530,830 | 13,286,970 | 12,085,033 | 90.95% | 6,485 | Discard |
| P5C2 | 20,486,302 | 16,468,422 | 13,941,479 | 12,255,904 | 87.91% | 5,107 | Discard |
| P5M1 | 19,619,732 | 15,668,424 | 12,861,530 | 11,169,688 | 86.85% | 8,646 | |
| P5M2 | 13,122,234 | 10,684,918 | 8,905,549 | 7,880,199 | 88.49% | 4,320 | Discard |
| P2S2C | 14,170,608 | 9,853,152 | 7,739,311 | 6,786,095 | 87.68% | 10,787 | |
| P2S3C | 15,838,436 | 13,011,848 | 10,886,651 | 10,035,120 | 92.18% | 3,213 | Discard |
| P2S1E | 18,630,710 | 11,671,876 | 6,581,134 | 5,469,753 | 83.11% | 7,934 | |
| P2S2M | 12,623,210 | 9,729,246 | 8,058,834 | 7,282,407 | 90.37% | 6,180 | |
| P2S3E | 5,884,824 | 4,363,930 | 2,340,440 | 2,108,293 | 90.08% | 4,737 | Discard |
| P3S4C | 15,204,128 | 11,651,710 | 9,749,853 | 9,000,435 | 92.31% | 7,228 | |
| P3S5C | 9,852,978 | 6,450,844 | 5,483,257 | 5,093,563 | 92.89% | 7,113 | |
| P3S2E | 13,250,672 | 9,012,308 | 7,705,896 | 7,054,340 | 91.54% | 3,376 | Discard |
| P3S4E | 11,546,022 | 7,638,676 | 6,447,749 | 5,880,059 | 91.20% | 7,444 | |
| P3S4M | 6,707,278 | 4,989,286 | 4,336,752 | 4,054,689 | 93.50% | 7,252 | |
| P3S5M | 13,655,400 | 9,380,232 | 7,933,731 | 7,068,156 | 89.09% | 6,370 | |
| P4S1C | 14,867,126 | 12,265,084 | 11,471,384 | 10,676,659 | 93.07% | 13,114 | |
| P4S2C | 12,633,350 | 9,891,910 | 8,953,036 | 8,496,549 | 94.90% | 12,299 | |
| P4S3C | 13,457,108 | 10,622,096 | 9,414,738 | 8,392,341 | 89.14% | 8,228 | |

**Supplementary Table 3** Summary of genomic DNA sequencing datasets.

| Sample | Raw Reads | Clean reads | Mapped Reads | Mapping ratio | CV | Note |
|---|---|---|---|---|---|---|
| P1S1C | 2,460,294 | 2,266,860 | 1,479,432 | 65.26% | 0.22 | |
| P1S3C | 1,982,770 | 1,824,536 | 1,199,801 | 65.76% | 0.25 | |
| P1S4C | 1,964,828 | 1,814,250 | 1,187,242 | 65.44% | 0.21 | |
| P1S1E | 1,998,404 | 1,852,388 | 1,206,034 | 65.11% | 0.20 | |
| P1S3G | 2,536,862 | 2,332,582 | 1,548,739 | 66.40% | 0.10 | |
| P1S4E | 2,653,418 | 2,487,224 | 1,638,529 | 65.88% | 0.30 | Discard |
| P1S4G | 2,887,628 | 2,647,064 | 1,449,046 | 54.74% | 0.23 | |
| P2S2C | 2,935,988 | 2,667,196 | 1,613,050 | 60.48% | 0.23 | |
| P2S3C | 3,174,466 | 2,996,854 | 682,293 | 22.77% | 0.39 | Discard |
| P2S1E | 2,962,818 | 2,779,362 | 1,633,143 | 58.76% | 0.15 | |
| P2S2M | 2,096,260 | 1,924,252 | 1,223,375 | 63.58% | 0.24 | |
| P2S3E | 3,701,746 | 3,440,192 | 2,158,518 | 62.74% | 0.14 | |
| P3S4C | 2,668,938 | 2,413,596 | 1,471,478 | 60.97% | 0.06 | |
| P3S5C | 2,880,934 | 2,618,020 | 1,593,564 | 60.87% | 0.12 | |
| P3S2E | 1,768,970 | 1,598,274 | 986,080 | 61.70% | 0.06 | |
| P3S4E | 2,620,464 | 2,146,864 | 555,029 | 25.85% | 0.27 | Discard |
| P3S4M | 2,391,802 | 2,176,122 | 1,314,536 | 60.41% | 0.06 | |
| P3S5M | 2,052,698 | 1,863,774 | 1,163,299 | 62.42% | 0.07 | |
| P4S1C | 3,333,178 | 3,103,906 | 1,934,962 | 62.34% | 0.20 | |
| P4S2C | 2,601,102 | 2,430,294 | 1,491,420 | 61.37% | 0.22 | |
| P4S3C | 3,715,784 | 3,533,988 | 2,262,832 | 64.03% | 0.39 | Discard |
| P4S1E | 2,995,878 | 2,790,760 | 1,750,622 | 62.73% | 0.10 | |
| P4S1M | 3,708,906 | 3,442,068 | 2,286,565 | 66.43% | 0.48 | Discard |
| P4S2E | 3,312,790 | 3,109,080 | 1,962,069 | 63.11% | 0.19 | |
| P4S2M | 2,563,728 | 2,036,234 | 1,048,489 | 51.49% | 0.20 | |
| P4S3E | 2,993,358 | 2,725,184 | 1,672,050 | 61.36% | 0.43 | Discard |
| P4S4M | 4,272,474 | 3,394,546 | 1,619,060 | 47.70% | 0.17 | |

Data files S1-SupExcel
